# Specific changes in sleep oscillations after blocking human metabotropic glutamate receptor 5 in the absence of altered memory function

**DOI:** 10.1101/2020.07.24.219865

**Authors:** Gordon B. Feld, Til Ole Bergmann, Marjan Alizadeh-Asfestani, Viola Stuke, Jan-Philipp Wriede, Surjo Soekadar, Jan Born

## Abstract

**Background:** Sleep consolidates declarative memory by repeated replay linked to the cardinal oscillations of NonREM sleep. However, there is so far little evidence of classical glutamatergic plasticity induced by this replay. Rather, we have previously reported that blocking NMDA or AMPA receptors does not affect sleep-dependent consolidation of declarative memory.

**Aims:** Investigate the role of metabotropic glutamate receptor 5 (mGluR5) on memory processing during sleep.

**Methods:** In two placebo-controlled within-subject cross-over experiments with 20 healthy humans each, we used fenobam to block mGluR5 during sleep. In Experiment I, participants learned word-pairs (declarative task) and a finger sequence (procedural task) in the evening, then received the drug and recall was tested in the next morning. To cover possible effects on synaptic renormalization processes during sleep, in Experiment II, participants learned new word-pairs in the morning after sleep.

**Results/Outcomes:** Surprisingly, fenobam neither reduced retention of memory across sleep nor new learning after sleep, although it severely altered sleep architecture and memory-relevant EEG oscillations. In NonREM sleep, fenobam suppressed 12-15 Hz spindles but augmented 2-4 Hz delta waves, whereas in REM sleep it suppressed 4-8 Hz theta and 16-22 Hz beta waves. Notably, under Fenobam NonREM spindles became more consistently phase-coupled to the slow oscillation.

**Conclusions/Interpretations:** Our findings indicate that mGluR5-related plasticity is not essential for memory processing during sleep, even though mGlurR5 are strongly implicated in the regulation of the cardinal sleep oscillations.

**Declaration of interest/Funding:** The authors have nothing to disclose and funders had no influence on the research presented here.

## Introduction

The memory enhancing function of sleep has been reported time and again (for a comprehensive review see: Klinzing et al., 2019; Rasch and Born, 2013, for current developments see: Feld and Born, 2017; Feld and Born, 2019, however also see Cordi and Rasch, 2021). Sleep’s role for memory relies on the repeated replay of memory traces that were encoded during prior wakefulness (Diekelmann and Born, 2010) and has been linked to the systems consolidation of hippocampus-dependent declarative memory (Dudai, 2004; Dudai et al., 2015). The hippocampo-neocortical dialog mediating this systems consolidation process is supported by the intricate interplay between slow oscillations (~0.75 Hz), sleep spindles (12-15 Hz) and sharp-wave/ripples (~100 Hz) that are observed during non-rapid-eyemovement (NonREM) sleep (Staresina et al., 2015; Molle and Born, 2011). While information flows from the sensory cortices to the hippocampus during wake, the direction of information flow is reversed during NonREM sleep (Mitra et al., 2016) and this reversal relies on low levels of acetylcholine during early slow wave sleep (SWS) rich NonREM sleep (Gais and Born, 2004; Rasch et al., 2006). Hippocampal replay as well as output from these neuron assemblies towards the neocortex is principally conveyed via glutamatergic neurons. Moreover, most researchers agree that replay and the associated neuronal oscillations are ideally suited to support systems consolidation by eliciting Hebbian plasticity in local synaptic circuitry (Khodagholy et al., 2017; Sadowski et al., 2016). Nevertheless, so far, our attempts to link sleep-dependent declarative memory consolidation to classical glutamatergic processes have largely failed (Feld et al., 2013). Here, we present our findings in pursuit of ascertaining the role of less well-characterized glutamatergic processes on sleep-dependent memory processes.

Classical glutamatergic N-methyl-D-aspartate (NMDA) receptor dependent long-term-potentiation (LTP) represents the most thoroughly investigated and, thus, prototypic form of neuronal plasticity in the mammalian brain (Huganir and Nicoll, 2013; Malenka and Bear, 2004). In brief, an incoming action potential elicits an excitatory post-synaptic potential at the post-synaptic neuron via sodium influx through the ion channel associated to the α-amino-3-hydroxy-5-methyl-4-isoxazolepropionic acid (AMPA) receptor. The NMDA receptor acts as a coincidence detector, inasmuch as, it is blocked by magnesium until the post-synaptic neuron is sufficiently depolarized by several incoming action potentials arriving within a certain time or space. In such a case, glutamate binding to the NMDA receptor allows the influx of calcium, finally, eliciting classical LTP, e.g., by enhancing signal transduction through AMPA receptor phosphorylation. In line with their essential role for brain plasticity, we could show that blocking NMDA or AMPA receptors during human sleep with ketamine or caroverine, respectively, completely abolished sleep-dependent performance gains for cortical memory (Gais et al., 2008). In contrast, applying the very same drugs during sleep did not impair the retention of declarative word pairs that relies on hippocampal areas for initial storage (Feld et al., 2013), although, the sleep-dependent consolidation of word-pairs was enhanced by the NMDA receptor co-agonist d-cycloserine.

The metabotropic glutamate receptor 5 (mGluR5) can be found throughout the hippocampus (Blumcke et al., 1996) and has been shown to contribute to NMDA receptor mediated LTP in the hippocampus (Xu et al., 2014; Ayala et al., 2009; Kroker et al., 2011). In humans, levels of mGluR5 were increased after sleep-deprivation (Hefti et al., 2013). A functional shift from the NMDA receptor to mGluR5 mediating enhanced calcium influx in the potentiated synapse offers a potential explanation for the above-mentioned negative findings after blocking of NMDA receptors during human sleep. In fact, activating mGluR5 has even been directly shown to reverse NMDA receptor blockade by ketamine (Lee et al., 2008).

mGluR5 has also been found to contribute to long-term-depression (LTD, Goh and Manahan-Vaughan, 2013; Ayala et al., 2009) – the opposite of LTP, inasmuch as, LTD reduces synaptic transmission by, e.g., de-phosphorylating AMPA receptors. This is of importance here because the replay-based systems consolidation process may indeed upscale cortical synapses to enhance a memory, but ripples may simultaneously weaken and downscale its hippocampal synapses (Norimoto et al., 2018). In addition, sleep exerts a global renormalizing effect on synaptic weights thereby erasing irrelevant memory traces and ensuring that the brain retains sustainability (Tononi and Cirelli, 2014; Feld and Born, 2017) which explains why the ability to learn new information is enhanced after sleep (Alizadeh Asfestani et al., 2018; Mander et al., 2011).

In the current research, we applied fenobam, a selective blocker of mGluR5 (Porter et al., 2005), during retention sleep in healthy humans. To investigate effects on sleep-dependent consolidation. In Experiment 1, participants learned word-pairs during the evening and then slept after receiving treatment. Retrieval was tested the next day. To additionally cover possible effects of fenobam on processes of synaptic downscaling and renormalisation, in Experiment 2, participants also received the drug before sleep after learning word-pairs. However, on the next morning their capability of learning new word-pairs was assessed. We expected that blockade of mGluR5 impairs sleep-dependent consolidation of the word-pairs as well as sleep-dependent improvements of new learning.

## Methods

### Participants

Forty participants completed the current study (n = 20 for each experiment). Participants were healthy, non-smoking, native German-speaking men (18–30 years), who reported no chronic physical or psychological illness in present or past. Participants underwent a routine health examination prior to participation to exclude any mental or physical disease, did not take any medication at the time of the experiments, and reported a normal sleep–wake cycle. The participants were instructed to get up at 07:00 on experimental days, and during these days not to take any naps and not to ingest alcohol or (after 13:00) caffeine-containing drinks. Before the experiment proper, participants took part in an adaption night under conditions of the experiment (i.e., including the placement of electrodes for polysomnographic recordings).The experiments were approved by the local ethics committee (Ethics Committee of the Medical Faculty at the University of Tübingen). Written informed consent was obtained from all participants before participation. Participation was compensated financially.

### Design and Procedures

Both experiments followed a double-blind, placebo-controlled, within-subject, balanced crossover design (see Figure 1A for an overview). Within each experiment, each participant underwent two identical sessions with the exception of receiving either fenobam or placebo, separated by at least 14 days. In the 1980s, fenobam was developed as a novel nonbenzodiazepine anxiolytic, but research was discontinued due to apparent neurological side effects (Pecknold et al., 1982). We chose fenobam due to its action as selective negative allosteric modulator at the mGluR 5 (Porter et al., 2005) and it was synthesised according to Good Laboratory Practice by Syncom (Groningen, Netherlands). Identical looking placebo and verum capsules were produced according to Good Manufacturing Practices by the Department of Pharmacy & Pharmacology at Slotervaart Hospital (Amsterdam, Netherlands). Each verum capsule contained 365 mg fenobam (plasma peak: 180 min, plasma half-life: 120 min, rough estimates according to Berry-Kravis et al., 2009). We chose this dose as it is the human equivalent dose of the dose that effectively blocked encoding in rats (Jacob et al., 2009). In this study, 30 mg/kg fenobam was more effective in blocking fear context conditioning than a dose of 10 mg/kg, but as effective as 100 mg/kg fenobam, making it the ideal choice after weighing the costs and benefits. Therefore, we divided the animal dose by 6.2, multiplied it by 75 kg and rounded to the nearest 5 mg increment, so that the dose could be prepared ahead of time at the pharmacy.

**Figure 1.**
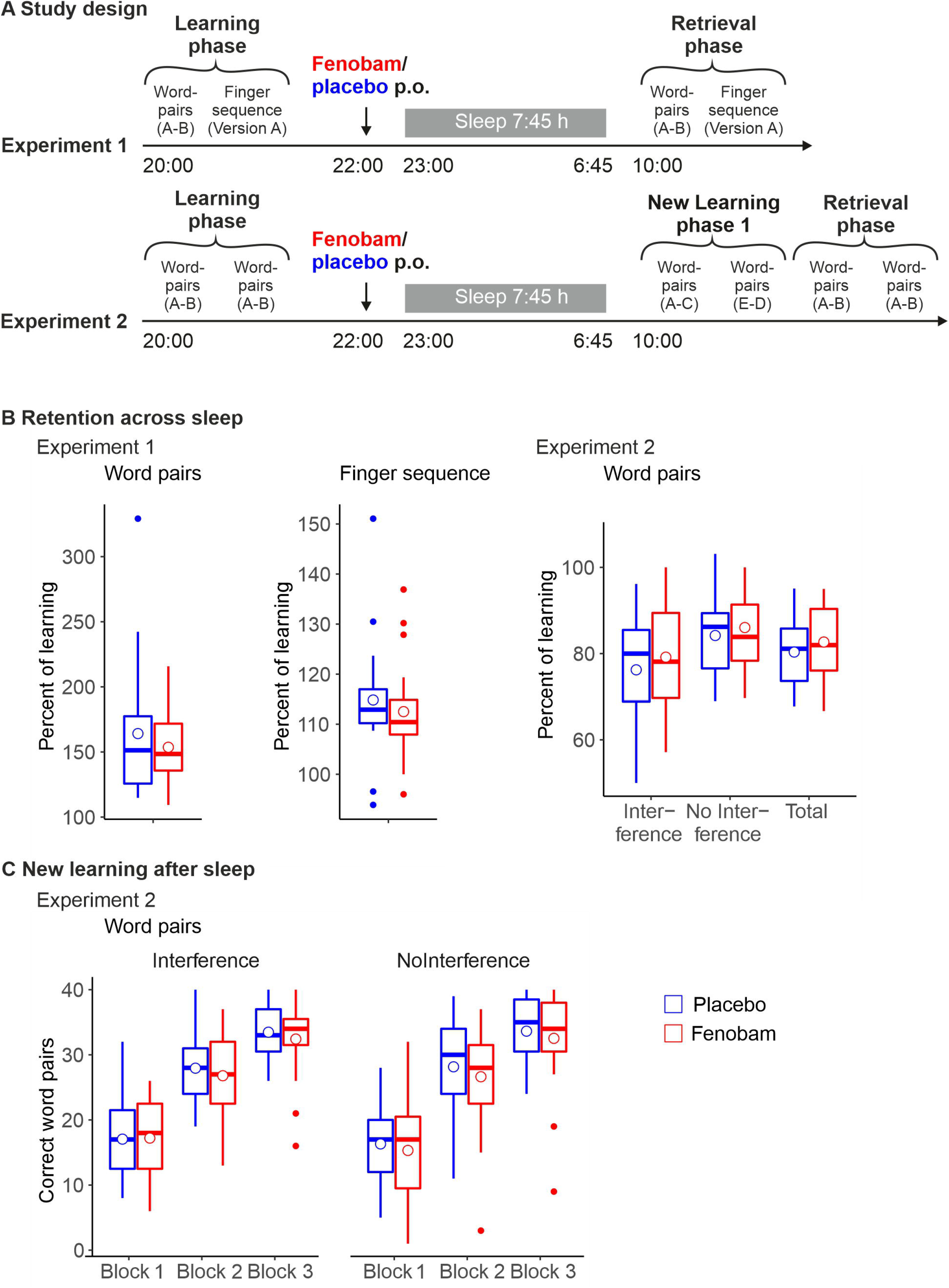
A) Time course of Experiment 1 and Experiment 2 (see Methods section for details). B) Correctly recalled word-pairs (A-B) during the Retrieval phase relative to the Learning phase in Experiment 1 (left panel). Correctly tapped finger sequences of the three blocks during the Retrieval phase relative to the last three blocks during the Learning phase in Experiment 1 (middle panel). Correctly recalled word-pairs during the Retrieval phase relative to the Learning phase in Experiment 2 for interference word-pairs (A-B), no interference word-pairs (A-B) and total (A-B) word-pairs (right panel). C) Mean (SEM) correctly recalled interfering (A-C) and noninterfering (E-D) word-pairs during the New Learning 1 phase in Experiment 2 (left and middle lower panel). Red and blue Box and whisker plots (thick line: median, hinge: 25% and 75% quartiles, whiskers: smallest/largest values no further than 1.5 * inter quartile range from the hinge) indicate fenobam and placebo, respectively.

#### Experiment 1

Participants arrived at the lab at 20:00 and were prepared for polysomnography. Next, they learned word-pairs and a finger sequence tapping task. Afterwards, vigilance, mood, adverse side effects and sleepiness were assessed and at 22:00 they received a capsule containing either fenobam or placebo. They watched animal documentaries until the electrodes were connected to the amplifier and at 23:00 the light was turned off. Participants were woken after 7:45-8:15 hours, preferentially from sleep stages 1 or 2 and sleepiness, adverse side effects as well as mood were assessed. Next, they were allowed to take a shower, they received a standardised breakfast and again watched animal documentaries to pass the time. At 10:00 retrieval was tested and additional control tasks were performed. Finally, vigilance, mood, adverse side effects and sleepiness as well as general retrieval performance were assessed.

#### Experiment 2

The timeline was identical to Experiment 1. However, we used a different protocol for word-pair learning in the evening and introduced learning of new word-pairs at 10:00 the next morning. In this experiment we did not assess finger sequence tapping.

### Memory Tasks

#### Word-pairs

The word-pair memory task was used to test hippocampus-dependent declarative memory. We decided to use more word-pairs than previously (Feld et al., 2013) to avoid ceiling effects and thus optimize the expected effect size (Feld et al., 2016a). In Experiment 1, participants were presented with 120 word-pairs (in three blocks of 40 word-pairs with a short break between each block) for 4 s each and with an inter stimulus interval (ISI) of 1 second during the Learning phase in the evening (procedure adapted from Ngo et al., 2013). Immediate recall was tested afterwards in a cued recall procedure, i.e., participants were shown the first word and had to respond by saying the second word. Irrespective of their performance they were shown the correct word as feedback for 2 s. In the next morning, during the Retrieval phase, delayed recall was tested using the same procedure but without feedback. Absolute differences between word pairs recalled during the Retrieval phase and during the Learning phase served as a measure of overnight retention.

In Experiment 2, participants were presented with 80 word-pairs (A-B in two blocks of 40) for 4 s each and with an ISI of 1 second during learning (procedure adapted from Feld et al., 2013 and Feld et al., 2016b). Immediate recall was tested using the same cued recall procedure as in Experiment 1. However, participants received no direct feedback and learning (3 s per stimulus) and immediate recall were repeated, if performance was below 60 %. During the New Learning phase before retrieval, participants learned 80 new word-pairs, half of which interfered with the original (A-B) word-pairs inasmuch as they shared the cue word (A-C), and half of which were entirely new (D-E). Word-pairs were learned in three blocks of learning followed by immediate cued recall. Word-pairs were presented for 4 s each during initial learning in the first block and for 3 s in the following block. Absolute values of word-pairs were used as a measure of new learning after sleep. During the subsequent Retrieval phase, recall of the original A-B word-pairs was tested again. Absolute differences between word-pairs recalled during the Retrieval phase and on the criterion trial during the Learning phase before sleep served as a measure of overnight retention.

#### Finger sequence tapping

In Experiment 1, the finger sequence tapping task was used to test procedural memory that is not critically depending on hippocampal function and has been described previously for having strong sleep-dependent effects (Nettersheim et al., 2015; Walker et al., 2003). Essentially, participants used their non-dominant hand to press a finger sequence (e.g., 4-1-3-2-4) on a keyboard as fast and as accurately as possible during twelve 30-second blocks interrupted by 30-second breaks. The numeric sequence was displayed on the screen at all times to keep working memory demands at a minimum. A key press resulted in a white asterisk appearing underneath the current element of the sequence.

Each 30-second block was scored for speed (number of correctly completed sequences) and errors. After each 30-second block, feedback was given about the number of correctly completed sequences and the error rate. During the Learning phase, participant trained on twelve 30-second blocks. The average score for the last three of these trials was used to indicate learning performance. During the Retrieval phase the next morning, participants were tested on another three blocks. The absolute difference between these was used as a score of retention performance. Afterwards, as a control for non-specific effects on learning a new sequence was learned for an additional 12 blocks.

### Polysomnography and Sleep Stage Analyses

The polysomnography was performed using Ag-AgCl electrodes connected to a Brain Amp DC 32-channel EEG-amplifier (Brain Products GmbH, Gilching) for the EEG channels and a Brain Amp ExG 8-channel bipolar amplifier (Brain Products GmbH, Gilching, Germany) that shared a ground connected to the forehead. The EEG was recorded from F3, Fz, F4, C3, Cz, C4, P3, Pz, P4 according to the international 10-20 system and referenced to coupled electrodes attached to the mastoids. Additionally, horizontal and vertical eye movements and muscle tone (from the chin) were recorded for standard polysomnography. The data were low-pass (80 Hz) and high-pass filtered (10 s time-constant) and sampled at 250 Hz. For sleep scoring the data were down-sampled to 200 Hz and additional offline low-pass (35 Hz) and high-pass (5 s time-constant) zero-phase Butterworth filters were applied. Sleep stage were scored according to Rechtschaffen and Kales (Rechtschaffen and Kales, 1968), using C3 and C4, by two experienced investigators who were blind to the assigned treatment Differences in scoring between the scorers were resolved by consulting a third experienced investigator. This sleep scoring approach, as still used in many scientific sleep labs around the world, was chosen to keep these data comparable to previous papers on the effect of the sleep on memory and augmented by additional sophisticated EEG analyses detailed below. In addition, we analysed sleep cycles across the night using the SleepCycles package in R (Blume and Cajochen, 2020).

### EEG analyses

During sleep scoring, it was evident that the drug influenced sleep EEG patterns severely enough to be identifiable by the naked eye during blind scoring (see supplementary file for example EEG traces). Therefore, in a subset of 24 participants (N = 11 from Experiment 1 and N = 13 from Experiment 2), for whom EEG signals in all 9 channels were of sufficient quality, more detailed EEG analysis were performed in MATLAB (v2016a, MathWorks, Natick, MA) using FieldTrip (Oostenveld et al., 2011) and CircStat (Berens, 2009) toolboxes as well as by applying custom MATLAB functions (see details in the sections below). Firstly, normalized, 1/f corrected power spectra during NonREM and REM sleep were compared between fenobam and placebo. Secondly, distinct oscillatory events were detected and characterized regarding their density, amplitude, and frequency. Finally, the phase-amplitude coupling (PAC) between slow oscillations (SO) and spindles was assessed, because of its known relevance for memory consolidation during sleep.

### Power spectral analysis

Power spectra were calculated using a Hanning-tapered Fast Fourier Transform (FFT) of non-overlapping artefact-free 4 s epochs (with a resolution of 0.25 Hz between 0.25 and 25 Hz), separately for NonREM and REM sleep. Note, we did not analyse the power spectrum during wake after lights out as after excluding epochs due to movement artefacts insufficient data remained. The analysis used the method of Irregular-Resampling Auto-Spectral Analysis (IRASA) (Wen and Liu, 2016), which has recently been used to successfully remove the 1/f noise from EEG power spectra obtained during sleep (Helfrich et al., 2018). In brief, IRASA resamples the time domain EEG data for each epoch using pairwise symmetric non-integer resampling factors (here: 1.1-1.9 in steps of 0.05 and corresponding factors 0.9-0.1) before calculating the FFT, resulting in multiple power spectra per epoch with artificially shifted peak frequencies for all oscillatory signals, while the 1/f power law noise component remains unchanged by resampling due to its fractal nature. Median-averaging of all power spectra per epoch then removes any frequency-shifted oscillatory signals and preserves only the fractal 1/f component, which in turn can be subtracted from the original power spectrum (of the non-resampled data) to obtain the 1/f-corrected power spectrum. We eventually calculated a normalized version of the 1/f-corrected power spectra by expressing the power in each frequency bin as the percentage change from the respective fractal power at that frequency. This approach elegantly normalizes oscillatory power per subject, channel, and frequency, thereby not only uncovering less pronounced oscillations that are usually hidden in 1/f noise, but also considerably reducing the large inter-subject variability in oscillatory power (see Supplementary Figure S1 for an illustrative example of the processing steps).

### Event detection analysis

To complement spectral power analyses, we detected and characterized discrete oscillatory events using detection algorithms established for SOs and spindles in NonREM sleep (Molle and Born, 2011; Mölle et al., 2002; Staresina et al., 2015). We used the latter in adapted form also to detect transient bursts of delta and theta oscillatory activity during NonREM sleep as well as theta, alpha, and beta oscillatory activity during REM sleep. Importantly, amplitude criteria for event detection were determined individually per night and channel. While a common absolute threshold or a common relative threshold across all nights enables confounding of oscillatory event detection by average oscillatory band power, relative thresholds determined per night control for potential power changes when analysing the density of discrete oscillatory events. In general, only artefact-free data from NonREM sleep stages S2-S4 or REM sleep were used for event detection, respectively, and data were always 50 Hz notch filtered to remove line noise. All bandpass-filters (see below) were zero-phase (two-pass) FIR filters with an order of 3 cycles of the respective low frequency cut-off.

For identifying SO events, data were bandpass-filtered 0.16-1.25 Hz, all zero-crossings were determined, and event duration of single waves was determined as time between two successive positive-to-negative zero-crossings. Events meeting the SO duration criteria (0.8-2 s corresponding to 0.5-1.25 Hz) were kept and their peak-to-peak amplitudes extracted. Events also meeting the SO amplitude criteria (greater than mean + 2 SD of the distribution of all extracted amplitudes) were then considered SOs, and epochs (−2.5 to +2.5 s) time-locked to the SO down-state negative peak in the filtered signal were extracted from the unfiltered raw signal for all events. For spindle detection, data were bandpass-filtered 12-15 Hz, and the root mean square (RMS) signal was calculated using a moving average of 200 ms. Whenever the signal exceeded the threshold of mean + 2 SD of the RMS signal (spindle amplitude criterion) for 0.5-3 s (spindle duration criterion) a spindle event was detected, and epochs (−2.5 to +2.5 s) time-locked to the maximum spindle trough in the filtered signal were extracted from the unfiltered raw signal for all events.

For the other oscillatory events, i.e., NonREM delta and theta as well as REM theta, alpha, and beta oscillatory events, detection procedures were analogous to the spindle detection pipeline, but with adapted bandpass-filter limits (delta: 2-4 Hz, theta: 4-8 Hz, alpha: 8-12 Hz, beta: 16-22 Hz) and event duration criteria (lower duration limit: 3 cycles of the centre frequency of the respective oscillatory band, e.g., for theta (2-4 Hz), at least 3 cycles at 6 Hz = 0.5 s; and no upper duration limit). Per subject, event density was calculated as number of events per minute, and average event amplitude was based on the respective maximal peak-to-peak amplitude of the filtered signal within the event limits. Individual peak frequency was derived from the normalized 1/f corrected power spectra as power maximum within the respective frequency band limits.

### SO-spindle phase-amplitude coupling (PAC) analysis

To assess the relationship between SO phase and spindle power we analysed SO-spindle phase-amplitude coupling (SO-spindle-PAC) as described in (Staresina et al., 2015). Both the phase of the SO bandpass-filtered signal and the phase of the spindle bandpass-filtered spindle power signal over time were extracted using the Hilbert transform. The spindle power modulation over time was estimated via a Hanning-tapered windowed FFT (FieldTrip’s ‘mtmconvol’ function) for frequencies between 12 and 15 Hz in steps of 1 Hz and a sliding window of 5 times the respective cycle length of a given frequency, moving in 10 ms steps. Frequency bins within the spindle range were averaged per time bin and upsampled to the EEG sampling rate of 250 Hz to align them with the SO phase estimates. The synchronization index between the two phase value time series (Cohen, 2008; Mormann et al., 2005), was then calculated for each SO epoch −1 to +1 s around the SO down-state. Averaging of phase-delays across SO events (CircStat toolbox) then provided for each participant the average SO phase angle at which spindle power was maximal (Figure 3D), as well as the consistency of this phase relationship across events, reflected by the respective mean vector length (Figure 3D). The deviation of individual SO-spindle-PAC phases from the group average indicated the consistency across participants (Figure 3E).

### Control Measures—Vigilance, Sleepiness, Mood Ratings, Word Fluency, Picture and Number Learning

Sleepiness and mood were assessed in the evening during the Learning phase as well as the next morning after awakening and during the Retrieval phase. Sleepiness was measured using the Stanford sleepiness scale (SSS; Hoddes et al., 1973). Mood was assessed using the multidimensional mood questionnaire (MDBF; Hinz et al., 2012). Additionally, during the Learning phase and the Retrieval phase vigilance was tested using a five minute version of the psychomotor vigilance task (PVT; Dinges et al., 1997) that required pressing a button as fast as possible whenever a bright millisecond clock, present on a dark computer screen, started counting upward. After the button press, this clock displayed the reaction time. Adverse side effects were measured by providing the most often encountered side effects under fenobam and offering an answer on a five point scale ranging from ‘not at all’ to ‘very much’ as well as a free text field.

Word fluency (Aschenbrenner et al., 2000) was tested at retrieval to control for effects on recall of long-term memories that have already been strongly consolidated. Participants produced as many words as possible prompted by a letter (p or m) and a category (hobby or occupation) cue for two minutes each. The sum of both versions was used a measure of general retrieval performance.

We additionally used a picture learning task and a number learning task at retrieval as non-verbal memory control tasks. In the picture learning task (that was only used in Experiment 1) participants viewed 120 pictures of landscapes and living room interiors for 1.5 s each (ISI 1-1.5 s). After a short break, immediate recognition was tested by presenting the learned pictures together with 120 new pictures. The discrimination index d-prime was used as measure of learning performance. In the number learning task (adapted from Feld et al., 2013) 16 three digit numbers were presented three times each for 2 s (500 ms ISI). A minute after presentation, participants were asked to freely recall the original numbers and free recall was used as a control measure of non-specific effects on learning after sleep. Afterwards, recognition of the numbers intermixed with 16 new three digit numbers was assessed, and d-prime was calculated as additional dependent measure.

### Statistical Analyses

Data from three participants were excluded due to poor sleep (two from Experiment 1 and one from Experiment 2). One additional participant was excluded from the sleep analyses in Experiment 2 due to missing data during one night and an additional participant was excluded from Experiment 1 as he had been allowed to sleep an hour longer in the fenobam condition by accident. In Experiment 1, the analyses used repeated measures ANOVAs with the factor Time point (reflecting performance at Learning and Retrieval phases), and Treatment (fenobam and placebo). In Experiment 2, we used factors for Interference (reflecting performance on the A-C interference word-pairs and the D-E no interference word-pairs), for Blocks (for learning block 1-3) and Treatment (fenobam and placebo). Post-hoc comparisons and other pairwise comparisons relied on dependent measures t-tests. Greenhouse-Geisser corrected degrees of freedom were used where appropriate. Cluster-based non-parametric permutation tests with correction for multiple comparisons across channels and frequency bins as implemented in FieldTrip (Maris and Oostenveld, 2007) were conducted to compare EEG power spectra under fenobam and placebo (Figures 3A and 4A). To allow post-hoc exploratory analyses of the time courses of spectral power changes across the night (despite the heterogeneous distribution of sleep stages preventing binning by time relative to drug intake or sleep onset), we divided all artifact-free epochs of each night and subject, separately for NonREM or REM sleep, into 10 equally-sized percentile bins, each containing 10% of consecutive data epochs (1-10%, 11-20%,…, 91-100%), and calculated the 1/f-corrected and normalized power values, separately for each percentile bin and drug condition (illustrated in Supplementary Figure S4). Power values extracted from each of the five significant channel x frequency clusters (i.e., NREM delta and spindle bands, and REM theta, alpha, and beta bands, cf. Figure 3+4) were then subjected to five separate two-way (2×10; Drug x Percentile) repeated-measures ANOVAs, and post-hoc one-way repeated-measures ANOVAs (separately per Drug condition) and post-hoc paired-t-tests (separately per Percentile) were conducted conditional on significant interaction effects (Figure 5). EEG measures from oscillatory event detection analyses (density, amplitude, frequency) were averaged across channels and subjected to two-sided paired t-tests contrasting fenobam and placebo conditions per frequency band (Figures 3C and 4C). For PAC analyses, Rayleigh tests were used to test for significant clustering of phase angles (deviation from uniform distribution) across subjects per condition, and two-sided paired t-tests were conducted to contrast mean vector length (Figure 3F) and average absolute deviation from group average (Figure 3G) between fenobam and placebo conditions.

## Results

### Memory tasks

In general, there were no significant effects of the treatment on measures of memory retention (all p > 0.14; see Figure 1B and Tables 1 and 2). Specifically, In Experiment 1, participants correctly recalled more word-pairs during the Retrieval phase than during the Learning phase (F_(1,16)_ = 137.52, p ≤ 0.001). None of the Treatment-related main or interaction effects were significant (p ≥ 0.61). Similarly, on the finger sequence tapping task, participants tapped more correct finger sequences during the Retrieval phase (F_(1,16)_ = 24.71, p ≤ 0.001), without any Treatment-related effects becoming apparent for correct sequences or the error rate (p ≥ 0.39). Participants increased the number of correctly tapped finger sequences during the for the control sequence learned after retrieval in Experiment 1 across the twelve blocks (F(3,48) = 21.10, p ≤ 0.001). But there were no further effects evident for the number of correct sequences or the error rate (p ≥ 0.27).

**Table 1:** Mean (SEM) for the memory tasks in Experiment 1.

|  | Fenobam |  | Placebo |  |
| --- | --- | --- | --- | --- |
| **Word-pairs** |  |  |  |  |
| Learning | 47.29 | (5.17) | 45.94 | (5.62) |
| Retrieval | 68.18 | (4.94) | 68.00 | (5.12) |
| Difference | 20.88 | (1.52) | 22.06 | (2.64) |
| Percent | 153.63 | (6.80) | 164.13 | (13.17) |
| **Finger sequence A** |  |  |  |  |
| **Correct sequences** |  |  |  |  |
| Learning | 20.63 | (0.95) | 20.69 | (1.29) |
| Retrieval | 23.33 | (1.39) | 23.82 | (1.69) |
| Difference | 2.71 | (0.64) | 3.14 | (0.63) |
| Percent | 112.51 | (2.64) | 114.83 | (3.10) |
| **Error rate** |  |  |  |  |
| Learning | 0.08 | (0.01) | 0.09 | (0.02) |
| Retrieval | 0.06 | (0.01) | 0.08 | (0.02) |
| Difference | -0.02 | (0.01) | -0.01 | (0.02) |
| **Finger sequence B** |  |  |  |  |
| **Correct sequences** |  |  |  |  |
| New learning 1-3 | 15.29 | (1.20) | 16.14 | (1.38) |
| New learning 4-6 | 18.29 | (1.26) | 19.20 | (1.29) |
| New learning 7-9 | 18.86 | (1.11) | 19.56 | (1.15) |
| New learning 10-12 | 19.64 | (1.37) | 19.35 | (1.21) |
| **Error rate** |  |  |  |  |
| New learning 1-3 | 0.11 | (0.01) | 0.13 | (0.03) |
| New learning 4-6 | 0.11 | (0.02) | 0.09 | (0.02) |
| New learning 7-9 | 0.10 | (0.01) | 0.08 | (0.01) |
| New learning 10-12 | 0.10 | (0.03) | 0.09 | (0.02) |
| **Numbers** |  |  |  |  |
| **Recognition** |  |  |  |  |
| D-prime | 1.45 | (0.20) | 1.71 | (0.19) |
| Hits | 0.72 | (0.04) | 0.72 | (0.04) |
| False alarms | 0.23 | (0.04) | 0.15 | (0.03) |
| **Free recall** |  |  |  |  |
| Absolute | 10.05 | (0.72) | 10.41 | (0.79) |
| Correct | 6.65 | (0.95) | 7.53 | (0.86) |
| **Pictures** |  |  |  |  |
| D-prime | 1.89 | (0.18) | 1.97 | (0.19) |
| Hits | 0.80 | (0.04) | 0.80 | (0.04) |
| False alarms | 0.20 | (0.03) | 0.19 | (0.03) |
| **Word generation** |  |  |  |  |
| Letter | 17.17 | (1.08) | 16.53 | (0.72) |
| Category | 21.82 | (1.22) | 20.00 | (0.84) |
| Total | 39.00 | (1.77) | 36.53 | (1.22) |

**Table 2:**
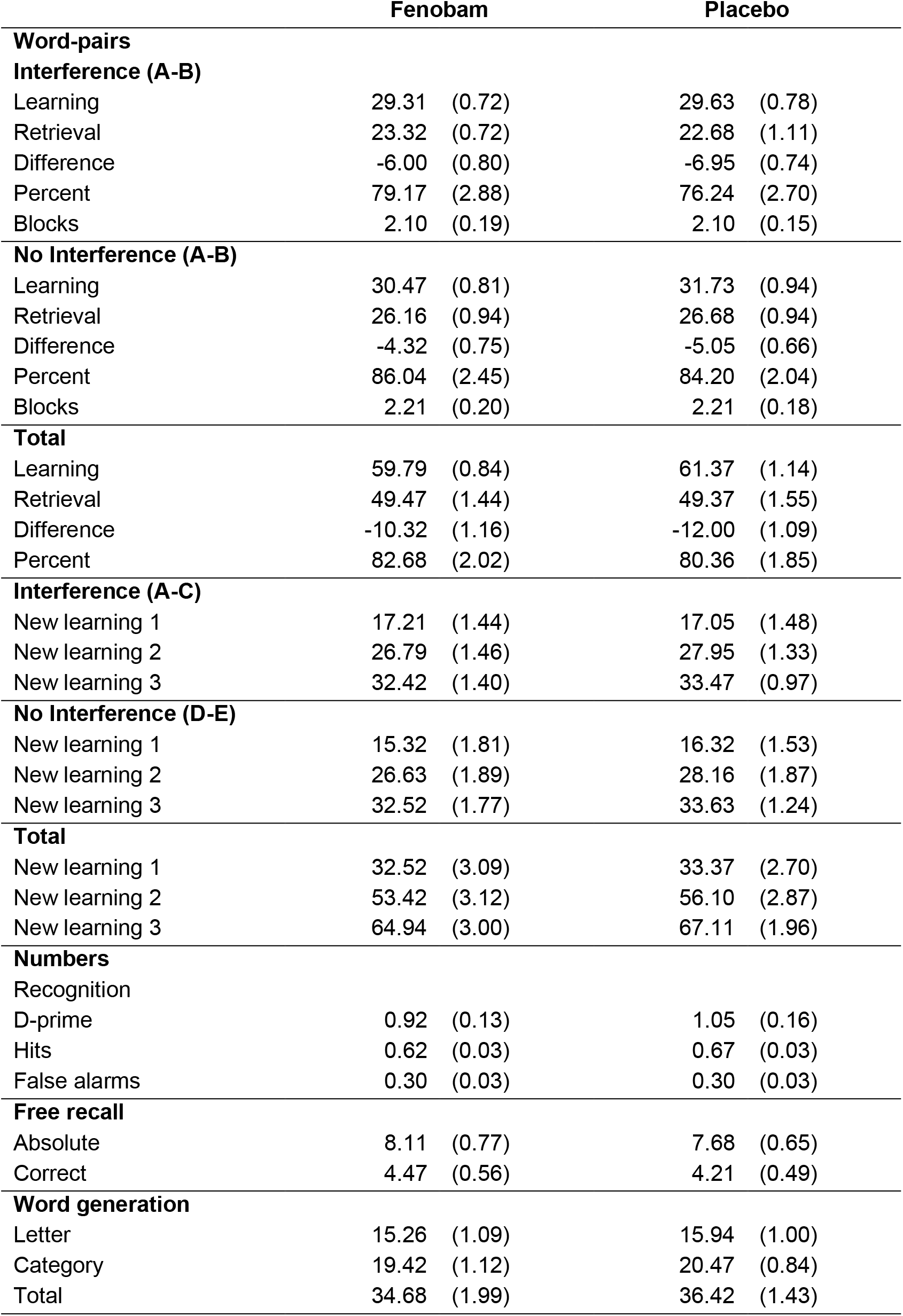
Mean (SEM) for the memory tasks in Experiment 2.

|  | Fenobam |  | Placebo |  |
| --- | --- | --- | --- | --- |
| Word-pairs |  |  |  |  |
| Interference (A-B) |  |  |  |  |
| Learning | 29.31 | (0.72) | 29.63 | (0.78) |
| Retrieval | 23.32 | (0.72) | 22.68 | (1.11) |
| Difference | -6.00 | (0.80) | -6.95 | (0.74) |
| Percent | 79.17 | (2.88) | 76.24 | (2.70) |
| Blocks | 2.10 | (0.19) | 2.10 | (0.15) |
| No Interference (A-B) |  |  |  |  |
| Learning | 30.47 | (0.81) | 31.73 | (0.94) |
| Retrieval | 26.16 | (0.94) | 26.68 | (0.94) |
| Difference | -4.32 | (0.75) | -5.05 | (0.66) |
| Percent | 86.04 | (2.45) | 84.20 | (2.04) |
| Blocks | 2.21 | (0.20) | 2.21 | (0.18) |
| Total |  |  |  |  |
| Learning | 59.79 | (0.84) | 61.37 | (1.14) |
| Retrieval | 49.47 | (1.44) | 49.37 | (1.55) |
| Difference | -10.32 | (1.16) | -12.00 | (1.09) |
| Percent | 82.68 | (2.02) | 80.36 | (1.85) |
| Interference (A-C) |  |  |  |  |
| New learning 1 | 17.21 | (1.44) | 17.05 | (1.48) |
| New learning 2 | 26.79 | (1.46) | 27.95 | (1.33) |
| New learning 3 | 32.42 | (1.40) | 33.47 | (0.97) |
| No Interference (D-E) |  |  |  |  |
| New learning 1 | 15.32 | (1.81) | 16.32 | (1.53) |
| New learning 2 | 26.63 | (1.89) | 28.16 | (1.87) |
| New learning 3 | 32.52 | (1.77) | 33.63 | (1.24) |
| Total |  |  |  |  |
| New learning 1 | 32.52 | (3.09) | 33.37 | (2.70) |
| New learning 2 | 53.42 | (3.12) | 56.10 | (2.87) |
| New learning 3 | 64.94 | (3.00) | 67.11 | (1.96) |
| Numbers |  |  |  |  |
| Recognition |  |  |  |  |
| D-prime | 0.92 | (0.13) | 1.05 | (0.16) |
| Hits | 0.62 | (0.03) | 0.67 | (0.03) |
| False alarms | 0.30 | (0.03) | 0.30 | (0.03) |
| Free recall |  |  |  |  |
| Absolute | 8.11 | (0.77) | 7.68 | (0.65) |
| Correct | 4.47 | (0.56) | 4.21 | (0.49) |
| Word generation |  |  |  |  |
| Letter | 15.26 | (1.09) | 15.94 | (1.00) |
| Category | 19.42 | (1.12) | 20.47 | (0.84) |
| Total | 34.68 | (1.99) | 36.42 | (1.43) |

In Experiment 2, participants recalled fewer word-pairs during the Retrieval phase than during the Learning phase (F_(1,18)_ = 126.90, p ≤ 0.001). Note, the generally worse recall in Experiment 2 than Experiment 1 is due to the procedure of Experiment 2 including, before the Retrieval phase, the New Learning of word-pairs half of which interfered (i.e., shared the same cue word) with the word-pairs learned during the Learning phase before sleep. Accordingly, there was also a main effect of interference (F_(1,18)_ = 7.28, p = 0.015) that was due to participants performing worse on retrieval of the interference word-pairs compared to the no-interference word-pairs during the Retrieval phase (time point x interference interaction: F_(1,18)_ = 6.92, p = 0.017). None of the Treatment-related effects was significant (p ≥ 0.14).

There were also no significant effects of Treatment on learning performance during the New Learning phase after sleep, in general (see Figure 1C and Tables 1 and 2 for descriptive statistics). In detail, during the New Learning of word pairs in Experiment 2 participants increased their performance across the three blocks (F_(2,36)_ = 355.56, p ≤ 0.001), but none of the other effects was significant (p ≥ 0.11).

### Sleep stages

Overall, fenobam severely impacted gross sleep architecture compared to placebo, when analysing time spent in each sleep stage relative to total sleep time (TST) during each night (Figure 2A and Table 3). The statistical analyses yielded the same results for absolute time in minutes per sleep stage, and consistent Treatment effects in both experiments. In Experiment 1, fenobam increased time spent in wakefulness and NonREM1 (wakefulness: t_(16)_ = −1.78, p = 0.095, NonREM1: t_(16)_ = −3.23, p = 0.005) in favour of NonREM 2, NonREM3 and REM sleep (NonREM2: t_(16)_ = 2.33, p = 0.033, NonREM3: t_(16)_ = 3.11, p = 0.007, p = 0.016, REM: t_(16)_ = 2.94, p = 0.010). NonREM4 and TST seemed unaffected by treatment (p ≥ 0.26). In Experiment 2, similarly, under fenobam participants spent more time in wakefulness and NonREM1 (wakefulness: t_(17)_ = −3.24, p = 0.005, NonREM1: t_(17)_ = −4.29, p ≤ 0.001) and less time in NonREM2 and NonREM3 (NonREM2: t_(17)_ = 3.57, p ≤ 0.001, NonREM3: t_(17)_ = 2.04, p = 0.057). Also, in Experiment 2 there was a trend towards participants sleeping less under fenobam than placebo (TST: t_(17)_ =, p = 0.089), but NonREM4 again stayed unaffected (p = 0.41).

**Figure 2.**
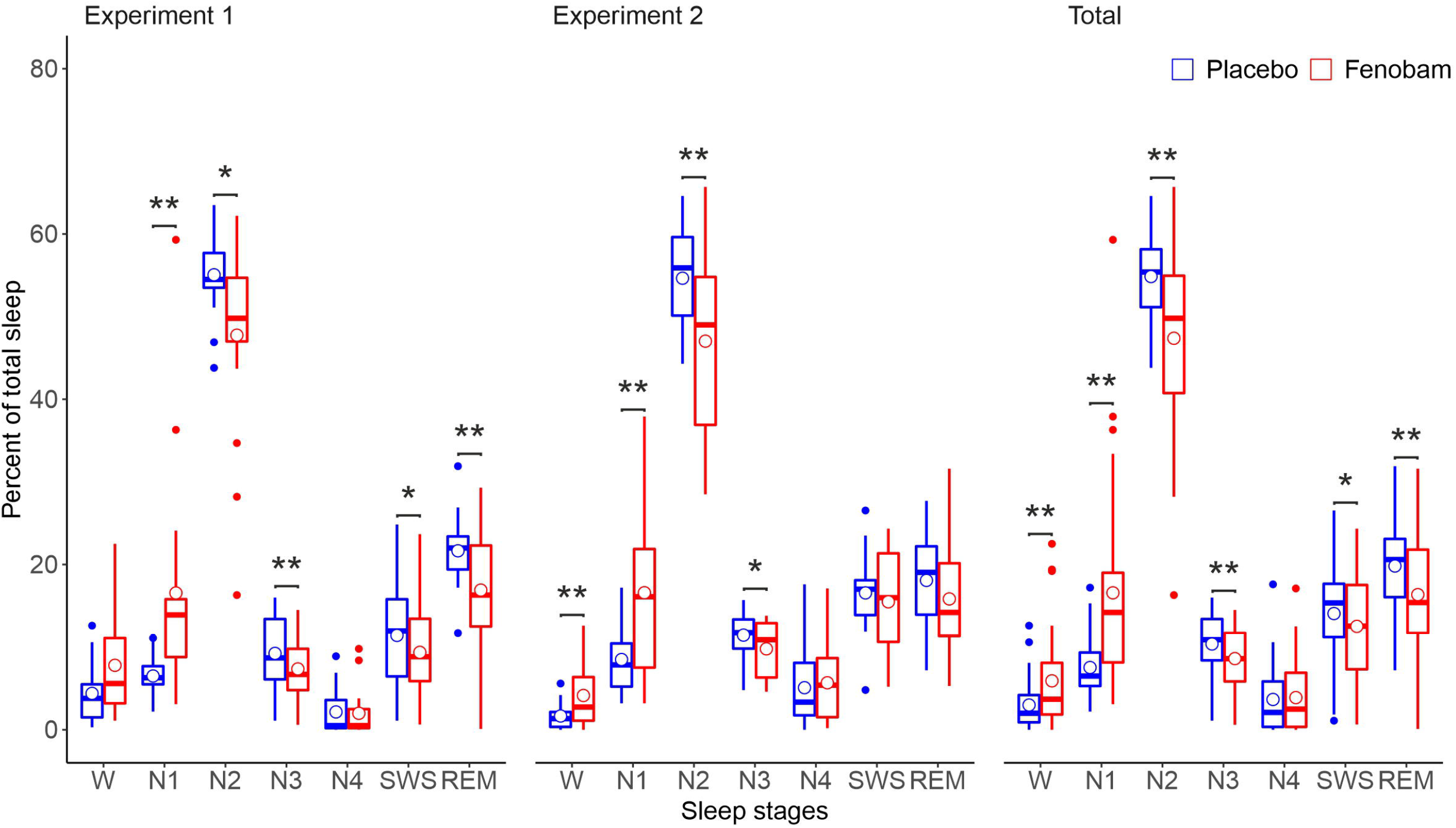
Mean (SEM) time spent in the different sleep stages (W – wake, N1-4 – NonREM1-4, SWS – slow wave sleep, REM – rapid eye movement sleep) relative to total sleep time for Experiment 1 (left panel), Experiment 2 (middle panel) and both experiments combined (right panel). Red and blue Box and whisker plots (thick line: median, hinge: 25% and 75% quartiles, whiskers: smallest/largest values no further than 1.5 * inter quartile range from the hinge) indicate fenobam and placebo respectively. (* p ≤ 0.05, ** p ≤ 0.01)

**Table 3:**
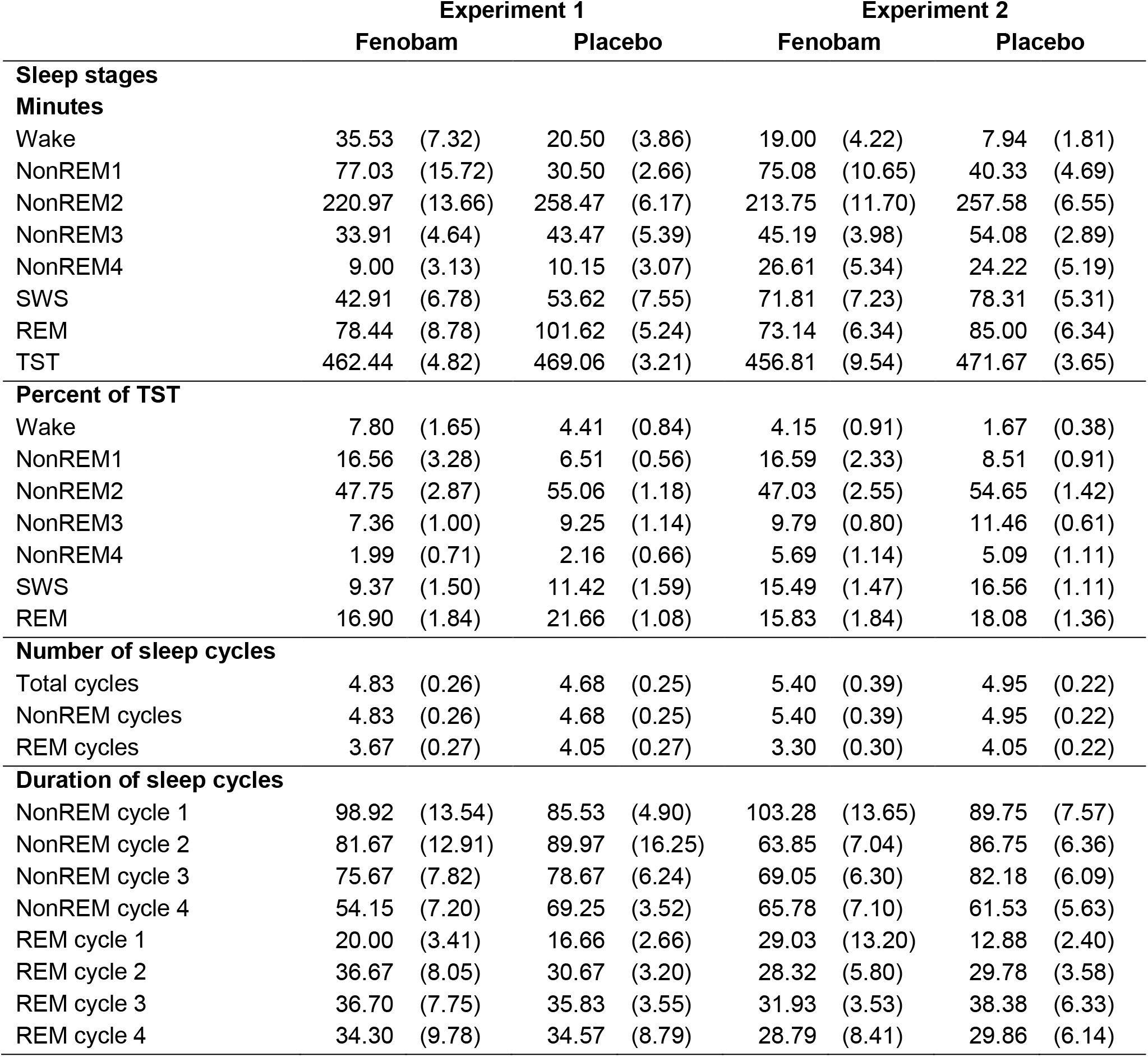
Mean (SEM) for the sleep stages and sleep cycles.

**Table 4:** Mean (SEM) for the the control measures. Note: Tiredness (low = tired), Positives affect (high = positive), Feeling calm (high = calm).

|  | Experiment 1 |  |  |  | Experiment 2 |  |  |  |
| --- | --- | --- | --- | --- | --- | --- | --- | --- |
|  | Fenobam |  | Placebo |  | Fenobam |  | Placebo |  |
| **Reaction speed (1/ms)** |  |  |  |  |  |  |  |  |
| Learning phase | 3.93 | (0.07) | 3.84 | (0.09) | 3.77 | (0.10) | 3.76 | (0.10) |
| Retrieval phase | 3.96 | (0.09) | 3.97 | (0.10) | 3.77 | (0.11) | 3.77 | (0.10) |
| **Sleepiness** |  |  |  |  |  |  |  |  |
| Learning phase | 3.94 | (0.30) | 3.82 | (0.23) | 2.89 | (0.21) | 3.16 | (0.21) |
| Awakening | 4.12 | (0.30) | 3.35 | (0.30) | 3.21 | (0.27) | 2.95 | (0.25) |
| Retrieval phase | 2.24 | (0.20) | 2.47 | (0.27) | 2.32 | (0.32) | 2.21 | (0.27) |
| **Adverse effects** |  |  |  |  |  |  |  |  |
| Learning phase | 2.29 | (0.44) | 2.47 | (0.30) | 1.73 | (0.35) | 2.16 | (0.49) |
| Awakening | 3.35 | (0.66) | 2.53 | (0.40) | 1.58 | (0.36) | 1.63 | (0.39) |
| Retrieval phase | 1.88 | (0.44) | 1.18 | (0.38) | 1.53 | (0.50) | 0.95 | (0.32) |
| **Tiredness** |  |  |  |  |  |  |  |  |
| Learning phase | 2.62 | 0.20 | 2.66 | 0.18 | 3.22 | 0.17 | 3.36 | 0.12 |
| Awakening | 2.75 | 0.20 | 3.04 | 0.21 | 3.14 | 0.18 | 3.37 | 0.17 |
| Retrieval phase | 3.56 | 0.20 | 3.59 | 0.25 | 3.65 | 0.25 | 3.78 | 0.23 |
| **Positive affect** |  |  |  |  |  |  |  |  |
| Learning phase | 3.63 | 0.13 | 3.42 | 0.20 | 4.13 | 0.08 | 4.48 | 0.08 |
| Awakening | 3.06 | 0.15 | 3.69 | 0.12 | 3.72 | 0.23 | 4.25 | 0.16 |
| Retrieval phase | 3.66 | 0.16 | 3.78 | 0.15 | 4.18 | 0.23 | 4.51 | 0.08 |
| **Feeling calm** |  |  |  |  |  |  |  |  |
| Learning phase | 3.83 | 0.16 | 3.62 | 0.21 | 3.88 | 0.18 | 4.18 | 0.16 |
| Awakening | 3.72 | 0.18 | 3.99 | 0.16 | 3.51 | 0.22 | 3.93 | 0.20 |
| Retrieval phase | 4.06 | 0.13 | 4.03 | 0.17 | 3.76 | 0.20 | 4.22 | 0.13 |
| **Medication guess** |  |  |  |  |  |  |  |  |
| Fenobam | 11 |  | 6 |  | 13 |  | 6 |  |
| Placebo | 5 |  | 12 |  | 6 |  | 13 |  |

Pooling sleep data from both experiments, to take advantage of the increased sample size, provided further evidence that fenobam increases wakefulness and NREM1 (wakefulness: t_(34)_ = −2.94, p = 0.006, NonREM1: t_(34)_ = −5.20, p ≤ 0.001), while decreasing NonREM2, NonREM3 and REM sleep (NonREM2: t_(34)_ = 4.04, p ≤ 0.001, NonREM3: t_(34)_ = 3.50, p ≤ 0.001, REM: t_(34)_ = 3.10, p = 0.004). It also provided more evidence for a general decrease in total sleep time under fenobam (TST: t_(34)_ = 2.16, p = 0.038). Even with this increased power to detect differences time spent in NonREM4 was not affected significantly (p = 0.59). A post-hoc analysis of the cycles of sleep during the night revealed that participants in the Fenobam condition had fewer sleep cycles including REM sleep (t_(33)_ = 2.31, p = 0.027).

### Power spectra, event detection, and phase-amplitude coupling (PAC) analyses

During NonREM stages 2-4 sleep fenobam (relative to placebo) suppressed spindle band (12-15 Hz) power but increased delta band (2-4 Hz) power in all channels (p_corr_ < 0.05; Figure 3A and Supplementary Figure S2). During REM sleep, theta band (4-8 Hz) and beta band (16-22 Hz) power were suppressed and alpha band (10-12 Hz) power was increased in all channels after fenobam (p_corr_ < 0.05; Figure 4A and Supplementary Figure S3). Thanks to the applied normalization and 1/f-correction procedure (Supplementary Figure S1), these findings cannot be attributed to changes in broadband power but have to originate from fenobam induced modulations of distinct oscillations in specific frequency bands. These changes lasted the entire night as demonstrated by post-hoc exploratory analyses of the spectral power time courses in these frequency bands (Figure 5). ANOVAs revealed significant main effects of Drug for all frequency bands (NREM Delta: F1,23 = 49.3, p = 0.0000004; NREM Spindle: F1,23 = 32.9, p = 0.000008; REM Theta: F1,23 = 24.2, p = 0.00006; REM Alpha: F1,23 = 13.5, p = 0.001; F1,23 = 23.6, REM Beta: p = 0.00007), and main effects (ME) of Percentile and Drug x percentile interactions (IA) only for NREM Spindle (ME: F9,207 = 2.3, p = 0.02; IA: F9,207 = 2.1, p = 0.03), REM Theta (ME trend only: F9,207 =1.8, p = 0.07; IA: F9,207 = 2.2, p =0.2), and REM Beta (ME: F9,207 = 5.7, p = 0.0000007; IA: F9,207 = 4.3, p = 0.00004). For frequency bands showing interactions, post-hoc tests (one-way ANOVAs across Percentile bins) revealed that these were driven by the fenobam-induced absence (all p > 0.2) of the modulation of oscillatory power otherwise observable across the night in the Placebo condition (NREM Spindle: F9,207 = 3.9, p = 0.0001; REM Theta: F9,207 = 3.0, p = 0.002; REM Beta: F9,207 = 10.4, p = 3×10-13), even though the difference between fenobam and placebo conditions remained for all percentiles tested (all p < 0.05).

**Figure 3.**
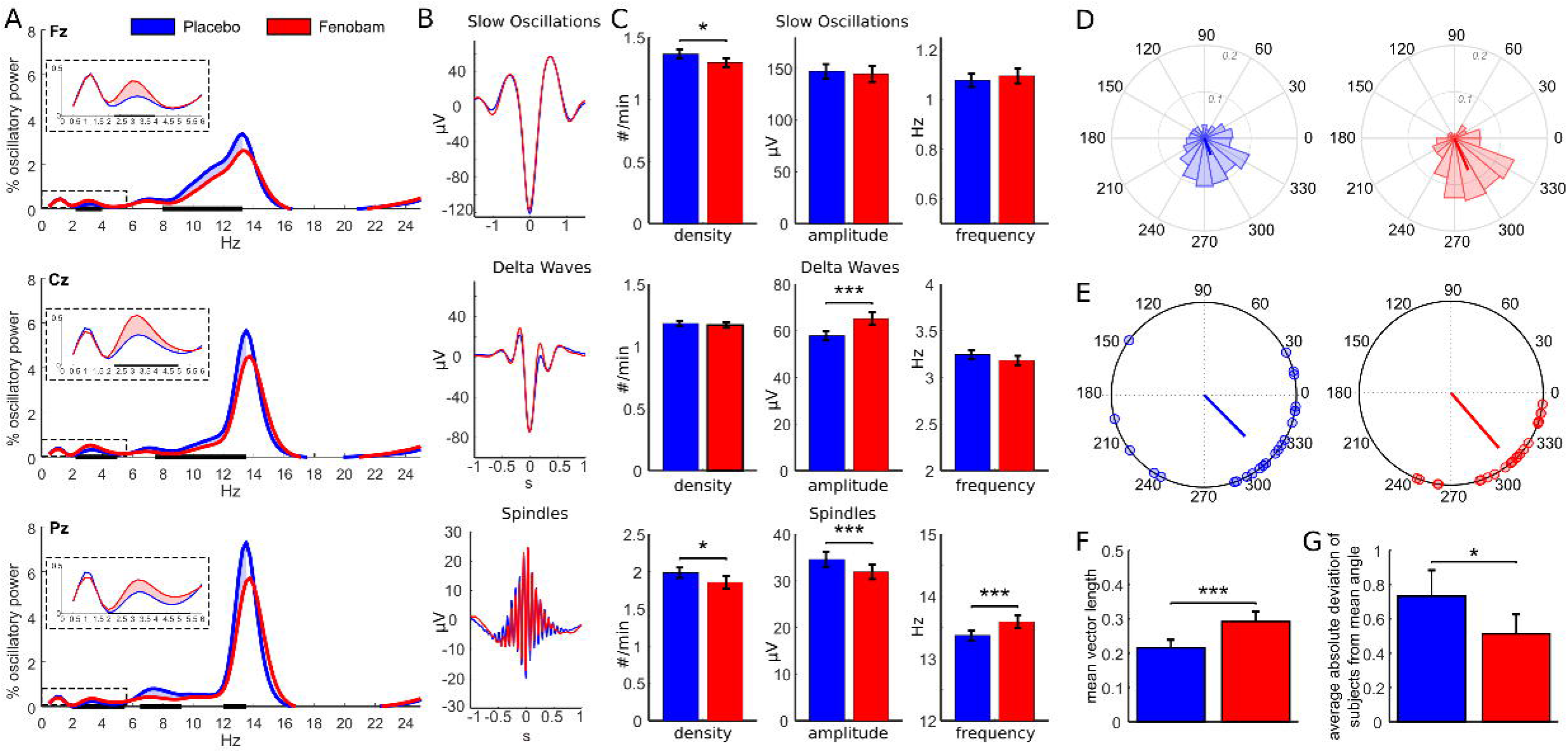
(A) Normalized 1/f-corrected power spectra during NonREM sleep (stages 2-4) reveal spindle (12-15 Hz) and theta (4-8 Hz) power decrease and delta power (2-4 Hz) increase after fenobam (red) compared to placebo (blue) intake (all 9 channels are shown in Supplementary Figure S2). Inserts provide zoom into outlined (dashed lines) areas to highlight delta power increases. Horizontal black bars and shaded area between lines indicate significant differences between fenobam and placebo (corrected for multiple comparisons across all electrodes and frequency bins using cluster-based permutation statistics). (B) Grand average time-locked waveforms for slow oscillations, delta waves, spindles resulting from event detection. (C) As revealed by event detection, under fenobam compared to placebo, slow oscillation and spindle density were reduced, spindle amplitude was decreased, delta wave amplitude increased, and spindles became slightly faster. Asterisks indicate significant differences at p < 0.05 (*), p < 0.01 (**), and p < 0.001 (***). (D) Angular histograms for a single representative participant illustrate slow oscillation to spindle phase-amplitude coupling (SO-spindle PAC) under fenobam (right) and placebo (left). (E) Preferred SO-spindle PAC phase (indicated by small circles on unitary circle) and their circular phase average across subjects (indicated by vector) reveal phase clustering across participants indicating spindle maxima during the slow oscillation down-to-up transition. Preferred SO-spindle PAC phase was comparable for fenobam (311°) and placebo (314°), but under fenobam SO-spindle PAC it was more clustered, both (F) within subjects (indicated by a larger mean vector length, p = 0.0007) and (G) across subjects (indicated by a smaller deviation of the subjects’ individual phase from the group mean, p = 0.015).

**Figure 4.**
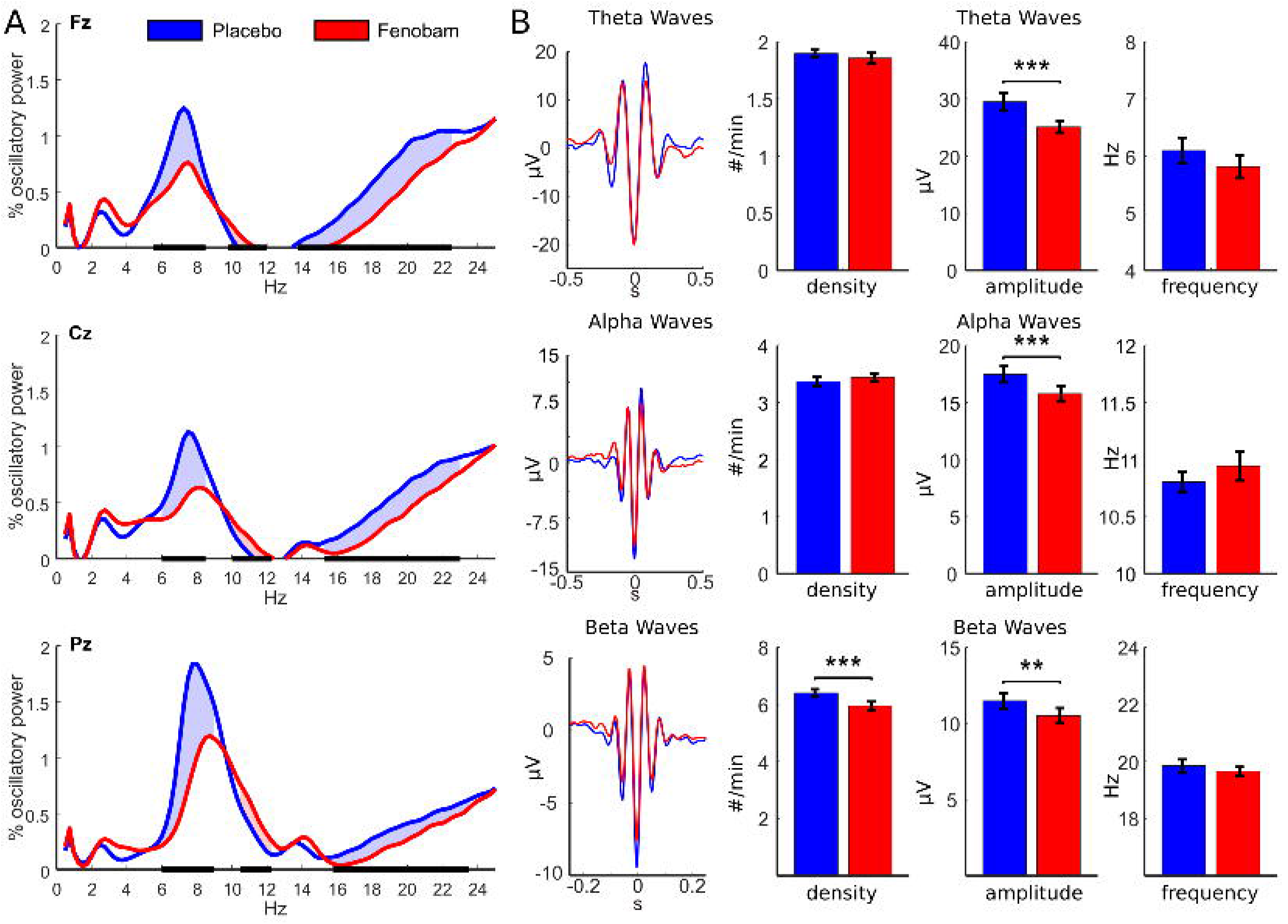
(A) Normalized 1/f-corrected power spectra during REM sleep reveal theta (4-8 Hz) and beta (16-22 Hz) power decreases and alpha power (10-12 Hz) increase (but no significant delta power increase) after fenobam (red) compared to placebo (blue) intake (all 9 channels are shown in Supplementary Figure S3). Horizontal black bars and shaded area between lines indicate significant differences between fenobam and placebo (corrected for multiple comparisons across all electrodes and frequency bins using cluster-based permutation statistics). (B) Grand average time-locked waveforms for delta, theta, and beta waves resulting from event detection. (C) As revealed by event detection, under fenobam compared to placebo, beta wave density was reduced, and theta, alpha, and beta wave amplitudes were decreased. Asterisks indicate significant differences at p < 0.05 (*), p < 0.01 (**), and p < 0.001 (***).

**Figure 5.**
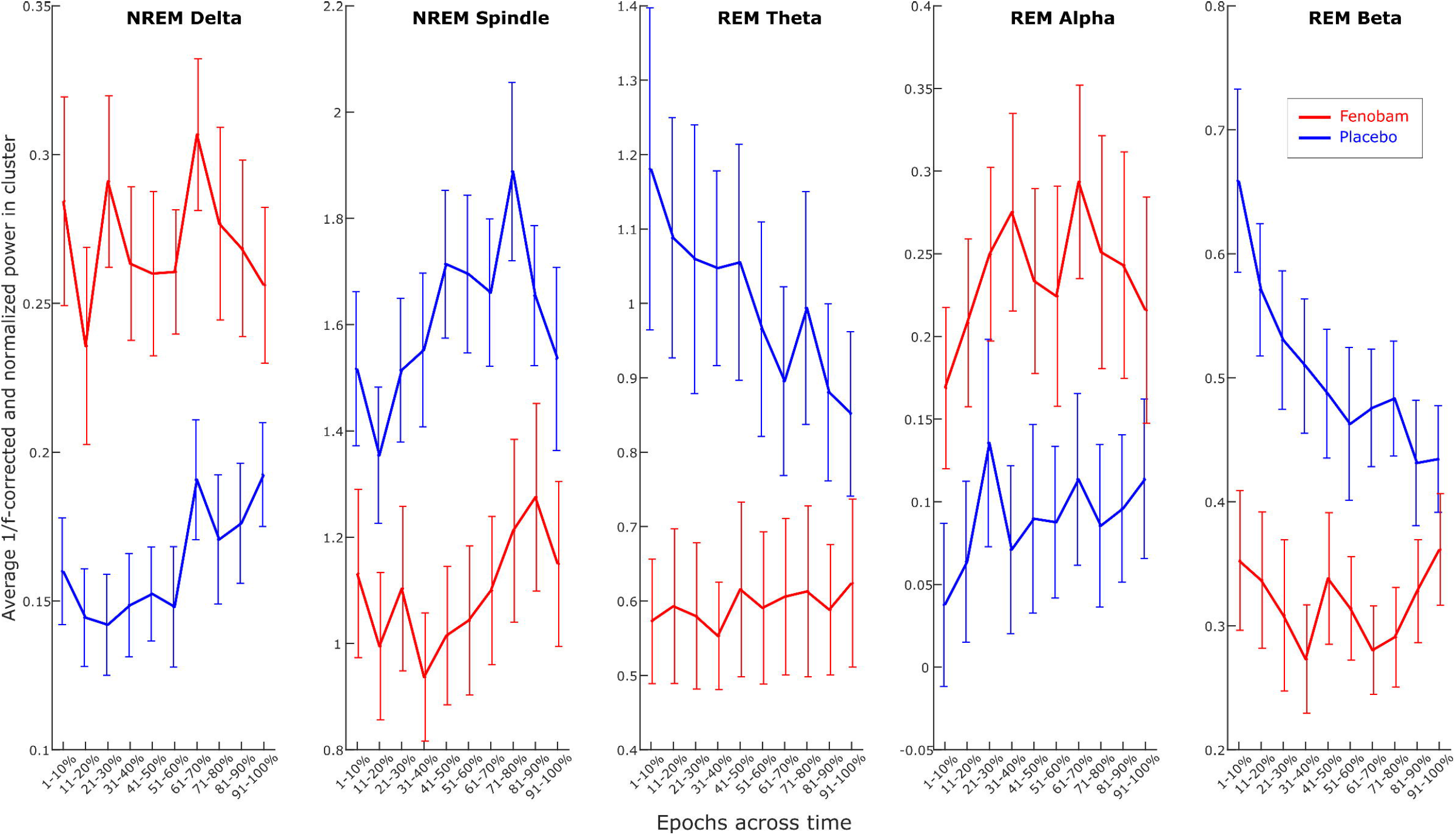
Averages (and SEM) of 1/f-corrected and normalized data in the frequency bands of interest, i.e., extracted from the significant channel x frequency clusters of the main analysis (cf. Figure 3+4), separately for NonREM delta and spindle bands as well as REM theta, alpha, and beta bands. Red and blue lines represent fenobam and placebo conditions, respectively. See Supplementary Figure 4 for full NonREM and REM power spectra of the same percentile bins. Note that power changes across the night are only significant for NonREM spindle, and REM theta and beta power in the placebo condition, while these modulations are suppressed following fenobam intake.

The findings of frequency-band specific power changes induced by fenobam were further corroborated by the oscillatory event detection analyses for these bands. During NonREM sleep (Figure 3B-C), fenobam indeed increased the amplitude of delta waves (t_23_ = – 4.21, p = 0.0003) while decreasing sleep spindle amplitude (t_23_ = 5.53, p < 0.0001). In addition, fenobam also reduced the density of distinct SOs (t_23_ = 2.26, p = 0.03) and spindle events (t_23_ = 2.50, p = 0.02) and increased spindle frequency (t_23_ = −3.93, p = 0.0008). During REM sleep (Figure 4 B-C), fenobam reduced the amplitude of theta (t_23_ = 3.78, p = 0.001), alpha (t_23_ = 6.27, p < 0.0001) and beta (t_23_ = 2.94, p = 0.007) waves as well as the density of beta waves (t_23_ = 3.99, p = 0.0006). Thus, only for the REM alpha oscillations there was an apparent mismatch between an increase in band power but a decrease in event amplitude, so that no unambiguous direction of modulation could be deduced for alpha oscillations.

We also tested whether the SO-spindle PAC was affected, which is thought to be most crucial for effective memory processing. Rayleigh tests confirmed significant clustering of spindle peaks at comparable phase-angles under fenobam (311°, z = 15.08, p < 0.00019) and placebo (314°, z = 9.62, p < 0.0000001). However, a larger mean vector length (t_23_ = 3.89, p < 0.001) and a smaller absolute deviation of individual preferred phase from the group angular mean was observed under fenobam than placebo (t_23_ = 2.63, p = 0.015), indicating that fenobam actually increased the precision and consistency of SO-spindle PAC both within and across subjects.

### Control measures

In Experiment 1, participants subjectively felt more sleepy in the morning after sleep under fenobam than under placebo according to the Stanford Sleepiness Scale (wakefulness: t_(16)_ = −2.34, p = 0.033), which was not the case in Experiment 2, at this time point, or at any other time points in both experiments (p ≥ 0.36). Objective measures of reaction speed (i.e., one divided by the individual average reaction time) on the psychomotor vigilance task did not show any treatment effects (p ≥ 0.17). The multidimensional mood questionnaire showed increased tiredness and reduced positive affect in the fenobam conditions directly after awakening in Experiment 1 (tiredness: t_(16)_ = 2.09, p ≤ 0.053, positive affect: t_(16)_ = 4.908, p ≤ 0.001). In Experiment 2, we found a baseline difference of reduced positive affect and feeling calm in the fenobam condition (positive affect: t_(17)_ = 2.70, p ≤ 0.015, feeling calm: t_(17)_ = 2.25, p ≤ 0.037), a trend towards lower positive affect in the fenobam condition directly after awakening (positive affect: t_(17)_ = 1,77, p = 0.094) and a reduced feeling of calmness after the Retrieval phase (feeling calm: t_(17)_ = 2.11, p = 0.049). All in all, any effects found in the control tasks were rather small und unsystematic, except for increased sleepiness at awakening in Experiment 1.

Concerning general retrieval performance measured by the word generation task during the Retrieval phase, there were no significant effects of treatment in Experiment 1 or Experiment 2 (p ≥ 0.18). When recognising numbers, participants made less false alarms in the placebo condition than in the fenobam condition in the New Learning phase of Experiment 1 (t_(16)_ = −2.14, p = 0.048), which was not replicated in the New Learning phase 2 of Experiment 2 (t_(18)_ ≤ 0.01, p ≥ 0.99). No other treatment effects were evident for the number learning task regarding recognition or free recall in Experiment 1 or Experiment 2 (p ≥ 0.19). There were also no significant treatment effects on recognising pictures in the New Learning phase of Experiment 1 (p ≥ 0.64).

There was no evidence of increased adverse side effects under fenobam in Experiment 1 or Experiment 2 at any point in time (p ≥ 0.17). However, in Experiment 2, there was a trend towards participants being able to guess whether they had received fenobam or placebo (Exact McNemar, p = 0.092) and this observation received further support when combining data from both experiments (Exact McNemar, p = 0.023).

## Discussion

Sleep benefits the formation of long-term memory by facilitating consolidation and new learning (Feld and Diekelmann, 2015). Sleep-dependent declarative memory consolidation has been shown to essentially rely on neuronal replay that is thought to lead to systems consolidation (Diekelmann and Born, 2010; Rasch and Born, 2013), whereas new learning after sleep has been proposed to benefit from synaptic renormalisation and presumed forgetting of irrelevant memories (Feld and Born, 2017; Tononi and Cirelli, 2014; Hardt et al., 2013). In the present study, we continue our efforts to identify the putative glutamatergic processes that are essential for these plastic processes at the synaptic level (Feld et al., 2013). Specifically, we hypothesised that blockade of mGluR5 during sleep by fenobam would impair replay-dependent synaptic consolidation processes. Moreover, given that these receptors might also be involved in synaptic depotentiation in hippocampal networks (Tadavarty et al., 2011) we explored whether the blocker would also impair new learning and encoding into hippocampal networks. To this end, participants learned word-pairs before sleep and retrieved them thereafter to measure retention in Experiment 1 and learned new word-pairs after sleep in Experiment 2, while receiving the selective mGluR5 blocker fenobam or placebo shortly before sleep. As expected, participants increased the number of recalled word-pairs across sleep in Experiment 1 as well as across learning blocks in Experiment 2. However, these processes were not impacted by fenobam. Interestingly, blocking mGluR5 had a strong effect on gross sleep architecture including sleep EEG oscillations closely linked to sleep-dependent memory processing. The mGluR5 blocker increased wakefulness and NonREM1 in favour of NonREM2, NonREM3 and REM sleep. More detailed analyses revealed that fenobam suppressed 12-15 Hz sleep spindles and augmented 2-4 Hz delta waves during NonREM, while suppressing mainly theta and beta waves during REM sleep. Notably, spindles became even more consistently coupled to slow oscillation phase, potentially compensating for their overall reduced number. Of note, increased delta activity above 2 Hz is compatible with reduced time spent in NREM2-4, as delta waves according to scoring guidelines must be below 2 Hz to signal deep sleep.

Glutamate is the most abundant excitatory neurotransmitter and glutamatergic LTP is the most intensively studied form of synaptic plasticity that leads to learning in neuronal systems (Malenka and Bear, 2004). To our surprise, we did not find evidence that blocking AMPA or NMDA receptors impairs sleep-dependent declarative memory consolidation (Feld et al., 2013). One possible explanation for this may be that the functional role of the NMDA receptor in mediating enhanced excitability shifts to other glutamatergic receptors during sleep. The mGluR5 is an ideal candidate for such a shift, as it is functionally coupled to the NMDA receptor (Chen et al., 2011). In line with this argument, mGluR5 receptor availability increases during sleep deprivation in healthy humans (Hefti et al., 2013), putatively acting as a tag for sleep-dependent consolidation. However, since we could not provide evidence for impaired sleep-dependent declarative memory consolidation by blocking mGluR5 such a shift seems unlikely to have acted here.

Not only synaptic potentiation but also depotentiation has been shown to rely on activation of mGuR5 (Chen et al., 2011; Tadavarty et al., 2011; Ayala et al., 2009). Similar to our findings regarding ketamine it is plausible that by blocking potentiation and depotentiation at the same time we received a null effect on the retention data. To answer this question we conducted Experiment 2, in which participants learned new word-pairs after sleep. Here, we also found no effect of mGluR5 blockade so this line of argument is refuted. Importantly, Experiment 2 also allowed us to conceptually replicate our null effect on retention in Experiment 1 with an interference paradigm, which has been shown to be sensitive to the memory enhancing effect of sleep (Ellenbogen et al., 2006).

Power is important when judging null effects. In this regard, the chosen sample size (n = 20 per experiment in a within-subject design) was a significant improvement compared to similar approaches of pharmacologically investigating sleep-dependent glutamatergic memory processes in humans (e.g., Feld et al., 2013; Gais et al., 2008) and constitutes a major experimental effort. We chose two procedures to measure declarative memory processing across sleep that have previously been shown to be highly susceptible to manipulations of sleep, yielding large effect sizes (Ngo et al., 2013; Ellenbogen et al., 2006; Feld et al., 2016a; Feld et al., 2016b). After excluding participants due to sleep problems we achieved reasonable power of 0.62 and 0.67 for Experiment 1 and Experiment 2 respectively (assuming a medium effect size of d = 0.5 and a two-sided type one error rate of α = 0.05, calculated in G*Power 3.1.7, Faul et al., 2007). When assessing both experiments together this yields the combined type two error rate of 12.54% for falsely retaining the null hypothesis that there is no medium to large effect of mGluR 5 blockade on memory processes in our experiments. To sum up, although smaller effects cannot reasonably be ruled out, we are confident that mGluR 5 mediated processes do not play a major role for sleep-dependent declarative memory processes.

Importantly, our finding that fenobam severely disrupted both REM and NonREM sleep and changed sleep-specific oscillatory activity, indicates that sufficient levels of fenobam arrived at the brain. Although effects of manipulating mGluR5 on sleep have been described in animal studies, they do not completely correspond to our findings in humans. In fact, some of the findings in rodents suggest the exact opposite relationship between mGluR5 activity and sleep (oscillatory activity) than what we observed in humans. Ahnaou and colleagues found that mGluR5 knock-out mice had a preserved circadian sleep-wake architecture, reduced REM sleep and reduced 1-4 Hz delta power during NonREM sleep, as well as reduced sleep drive and SWA (0.5-4 Hz) after sleep deprivation (Ahnaou et al., 2015b). In rats, mGluR5 negative allosteric modulators also suppressed REM sleep, but increased and consolidated NonREM sleep (Ahnaou et al., 2015a; Cavas et al., 2013), whereas mGluR5 positive allosteric modulators increased waking, and decreased both REM and NonREM sleep (Ahnaou et al., 2015a). Others reported mGluR5 to be wake promoting in general (Gilmour et al., 2013).

In line with a sleep regulatory role of mGluR 5, a recent positron emission tomography (PET) study in humans, using a highly selective, non-competitive radio-ligand, showed increased mGluR5 availability after sleep loss (Holst et al., 2017). This increase was correlated to behavioural and EEG biomarkers of elevated sleep need, with mGluR5 availability being positively related to NonREM slow wave activity (0.5-4.5 Hz), both at baseline and after sleep deprivation. In this sense, mGluR5 mediated activity seems to support homeostatic increases in SWA in humans, which is reconcilable with the reduced time spent in SWS after blocking these receptors with fenobam in our study. Note, that we also found the density of SOs (0.5-1.25 Hz) to be reduced, but the amplitude of delta waves (2-4 Hz to be increased under fenobam. While corroborating the different nature of these two oscillations (Steriade, 2006), it complicates a direct comparison with studies for which SO and delta were lumped together as slow wave activity (SWA; 0.5-4.5 Hz). We speculate that the spectrally circumscribed increase in delta power, resulted from an increase in thalamically, as opposed to cortically, generated slow waves (Steriade, 2006). While it is very plausible that metabotropic glutamate receptors in the thalamus and cortico-thalamic feedback mechanisms are involved in the observed effects of fenobam on spindle decreases and delta increases, the direction of the effect is again opposed to what would have been predicted from previous animal work. Corticothalamic activation of metabotropic glutamate receptors in the thalamus of guinea pigs induced a slow depolarization via reduced potassium currents, shifting the burst firing mode of thalamic relay cells, characteristic for SWS, to the single spiking mode dominating wakefulness (McCormick and von Krosigk, 1992), and corticothalamic feedback has specifically been shown to be capable of flipping thalamic activity from spindle bursts to ~3 Hz delta bursts (even though metabotropic glutamate receptors were not considered in those studies, Bal et al., 2000; Destexhe et al., 1998). It therefore remains unclear why in humans blocking of mGluR5 (instead of its activation) caused exactly such a shift from spindle to delta waves. The present findings thus corroborate the idea that mGluR5 are significantly involved in the regulation of sleep and that they exert their influence on sleep specific oscillatory activity via corticothalamic-feedback loops. However, the direction of the effects observed in humans and rodents is not easily reconciled, and specific studies aimed at disentangling the precise mechanisms at play are needed. Notably, the current study did not vary the dose of fenobam and it is conceivable that at other dosing effects of the drug in humans would align more clearly with those found in animals. Also, the relative amount of SWS in the placebo condition was on the low side, which may signal additional nocebo effects of drug intake.

Irrespective of the precise pharmacological and neurophysiological underpinnings of the observed spindle suppression, the question remains why a significant decrease in spindle amplitude and density did not affect memory consolidation. Recent work has demonstrated that memory consolidation relies in particular on spindles occurring during SO up-states, reflected by SO-spindle-PAC (Bergmann and Born, 2018; Niethard et al., 2018; Helfrich et al., 2018; Latchoumane et al., 2017). Notably, SO-spindle-PAC was not only preserved under fenobam, but became even more pronounced, both within and across subjects. Unlike their overall reduction in power and density, the increased synchronization of spindles to SO upstates is well in line with a fenobam-induced antagonism of mGluR5 receptors in the thalamus, which is supposed to facilitate the SWS-like burst firing mode of thalamocortical relay cells (McCormick and von Krosigk, 1992).

In conclusion, our experiments constitute yet another failure to find essential contributions of glutamatergic processes to sleep-dependent memory formation that adds to our finding that AMPA and NMDA receptor blockade does not impair sleep-dependent declarative memory consolidation (Feld et al., 2013). This remains an especially unexpected finding considering the importance of glutamatergic contributions to neurotransmission, neuroplasticity and learning, which makes it important to emphasize that for example blockade of NMDA-receptor subunits (e.g., NR2B) or simultaneous blockade of NMDA and mGlu 5 receptors could be a more promising approach. However, this can only be done in animal models. While it may seem that our work is marginalising glutamate, quite to the contrary, it is possible that glutamatergic contributions to sleep-dependent memory processes are so important for an organism’s survival that there exist several redundant signalling pathways. In addition, our findings of sleep disruption, reduced sleep-spindle activity and increases in un-physiological high delta underline the importance of mGluR5 for sleep regulation. Considering current efforts to develop mGluR5 related drugs (Pillai and Tipre, 2016), it will be highly important to characterise sleep disturbance (or improvement) related to these drugs as part of their neuropsychopharmacological profile.

## References

Ahnaou A, Langlois X, Steckler T, et al. (2015a) Negative versus positive allosteric modulation of metabotropic glutamate receptors (mGluR5): indices for potential pro-cognitive drug properties based on EEG network oscillations and sleep-wake organization in rats. Psychopharmacology (Berl) 232(6): 1107–1122.

Ahnaou A, Raeymaekers L, Steckler T, et al. (2015b) Relevance of the metabotropic glutamate receptor (mGluR5) in the regulation of NREM-REM sleep cycle and homeostasis: evidence from mGluR5 (-/-) mice. Behav Brain Res 282: 218–226.

Alizadeh Asfestani M, Braganza E, Schwidetzky J, et al. (2018) Overnight memory consolidation facilitates rather than interferes with new learning of similar materials-a study probing NMDA receptors. Neuropsychopharmacology 43(11): 2292–2298.

Aschenbrenner S, Tucha O and Lange KW (2000) Regensburger Wortflüssigkeits-Test: RWT. Hogrefe, Verlag für Psychologie.

Ayala JE, Chen Y, Banko JL, et al. (2009) mGluR5 positive allosteric modulators facilitate both hippocampal LTP and LTD and enhance spatial learning. Neuropsychopharmacology 34(9): 2057–2071.

Bal T, Debay D and Destexhe A (2000) Cortical feedback controls the frequency and synchrony of oscillations in the visual thalamus. J Neurosci 20(19): 7478–7488.

Berens PJJSS (2009) CircStat: a MATLAB toolbox for circular statistics. 31(10): 1–21.

Bergmann TO and Born J (2018) Phase-Amplitude Coupling: A General Mechanism for Memory Processing and Synaptic Plasticity? Neuron 97(1): 10–13.

Berry-Kravis E, Hessl D, Coffey S, et al. (2009) A pilot open label, single dose trial of fenobam in adults with fragile X syndrome. J Med Genet 46(4): 266–271.

Blumcke I, Behle K, Malitschek B, et al. (1996) Immunohistochemical distribution of metabotropic glutamate receptor subtypes mGluR1b, mGluR2/3, mGluR4a and mGluR5 in human hippocampus. Brain Res 736(1-2): 217–226.

Blume C and Cajochen C. (2020). ‘SleepCycles’ package for R – A free software tool for the detection of sleep cycles from sleep staging. Open Science Framework (OSF). doi: https://doi.org/10.31219/osf.io/r2q8v

Cavas M, Scesa G and Navarro JF (2013) Effects of MPEP, a selective metabotropic glutamate mGlu5 ligand, on sleep and wakefulness in the rat. Prog Neuropsychopharmacol Biol Psychiatry 40: 18–25.

Chen HH, Liao PF and Chan MH (2011) mGluR5 positive modulators both potentiate activation and restore inhibition in NMDA receptors by PKC dependent pathway. J Biomed Sci 18: 19.

Cohen MX (2008) Assessing transient cross-frequency coupling in EEG data. J Neurosci Methods 168(2): 494–499.

Cordi MJ and Rasch B (2021) How robust are sleep-mediated memory benefits? Curr Opin Neurobiol 67.

Destexhe A, Contreras D and Steriade M (1998) Mechanisms underlying the synchronizing action of corticothalamic feedback through inhibition of thalamic relay cells. J Neurophysiol 79(2): 999–1016.

Diekelmann S and Born J (2010) The memory function of sleep. Nat Rev Neurosci 11(2): 114–126.

Dinges DF, Pack F, Williams K, et al. (1997) Cumulative sleepiness, mood disturbance, and psychomotor vigilance performance decrements during a week of sleep restricted to 4-5 hours per night. Sleep 20(4): 267–277.

Dudai Y (2004) The neurobiology of consolidations, or, how stable is the engram? Annu Rev Psychol 55: 51–86.

Dudai Y, Karni A and Born J (2015) The Consolidation and Transformation of Memory. Neuron 88(1): 20–32.

Ellenbogen JM, Hulbert JC, Stickgold R, et al. (2006) Interfering with theories of sleep and memory: sleep, declarative memory, and associative interference. Curr Biol 16(13): 1290–1294.

Faul F, Erdfelder E, Lang AG, et al. (2007) G*Power 3: a flexible statistical power analysis program for the social, behavioral, and biomedical sciences. Behav Res Methods 39(2): 175–191.

Feld GB and Born J (2017) Sculpting memory during sleep: concurrent consolidation and forgetting. Curr Opin Neurobiol 44: 20–27.

Feld GB and Born J (2020) Neurochemical mechanisms for memory processing during sleep: basic findings in humans and neuropsychiatric implications. Neuropsychopharmacology 45: 31–44.

Feld GB and Diekelmann S (2015) Sleep smart-optimizing sleep for declarative learning and memory. Front Psychol 6: 622.

Feld GB, Lange T, Gais S, et al. (2013) Sleep-dependent declarative memory consolidation-unaffected after blocking NMDA or AMPA receptors but enhanced by NMDA coagonist D-cycloserine. Neuropsychopharmacology 38(13): 2688–2697.

Feld GB, Weis PP and Born J (2016a) The Limited Capacity of Sleep-Dependent Memory Consolidation. Front Psychol 7.

Feld GB, Wilhem I, Benedict C, et al. (2016b) Central Nervous Insulin Signaling in Sleep-Associated Memory Formation and Neuroendocrine Regulation. Neuropsychopharmacology 41(6): 1540–1550.

Gais S and Born J (2004) Low acetylcholine during slow-wave sleep is critical for declarative memory consolidation. Proc Natl Acad Sci U S A 101(7): 2140–2144.

Gais S, Rasch B, Wagner U, et al. (2008) Visual-procedural memory consolidation during sleep blocked by glutamatergic receptor antagonists. J Neurosci 28(21): 5513–5518.

Gilmour G, Broad LM, Wafford KA, et al. (2013) In vitro characterisation of the novel positive allosteric modulators of the mGlu(5) receptor, LSN2463359 and LSN2814617, and their effects on sleep architecture and operant responding in the rat. Neuropharmacology 64: 224–239.

Goh JJ and Manahan-Vaughan D (2013) Endogenous hippocampal LTD that is enabled by spatial object recognition requires activation of NMDA receptors and the metabotropic glutamate receptor, mGlu5. Hippocampus 23(2): 129–138.

Hardt O, Nader K and Nadel L (2013) Decay happens: the role of active forgetting in memory. Trends Cogn Sci 17(3): 111–120.

Hefti K, Holst SC, Sovago J, et al. (2013) Increased metabotropic glutamate receptor subtype 5 availability in human brain after one night without sleep. Biol Psychiatry 73(2): 161–168.

Helfrich RF, Mander BA, Jagust WJ, et al. (2018) Old Brains Come Uncoupled in Sleep: Slow Wave-Spindle Synchrony, Brain Atrophy, and Forgetting. Neuron 97(1): 221–230 e224.

Hinz A, Daig I, Petrowski K, et al. (2012) [Mood in the German population: norms of the Multidimensional Mood Questionnaire MDBF]. Psychother Psychosom Med Psychol 62(2): 52–57.

Hoddes E, Zarcone V, Smythe H, et al. (1973) Quantification of sleepiness: a new approach. Psychophysiology 10(4): 431–436.

Holst SC, Sousek A, Hefti K, et al. (2017) Cerebral mGluR5 availability contributes to elevated sleep need and behavioral adjustment after sleep deprivation. Elife 6.

Huganir RL and Nicoll RA (2013) AMPARs and synaptic plasticity: the last 25 years. Neuron 80(3): 704–717.

Jacob W, Gravius A, Pietraszek M, et al. (2009) The anxiolytic and analgesic properties of fenobam, a potent mGlu5 receptor antagonist, in relation to the impairment of learning. Neuropharmacology 57(2): 97–108.

Khodagholy D, Gelinas JN and Buzsaki G (2017) Learning-enhanced coupling between ripple oscillations in association cortices and hippocampus. Science 358(6361): 369–372.

Klinzing JG, Niethard N and Born J (2019) Mechanisms of systems memory consolidation during sleep. Nat Neurosci 22(10): 1598–1610.

Kroker KS, Rast G and Rosenbrock H (2011) Differential effect of the mGlu5 receptor positive allosteric modulator ADX-47273 on early and late hippocampal LTP. Neuropharmacology 61(4): 707–714.

Latchoumane CV, Ngo HV, Born J, et al. (2017) Thalamic Spindles Promote Memory Formation during Sleep through Triple Phase-Locking of Cortical, Thalamic, and Hippocampal Rhythms. Neuron 95(2): 424–435 e426.

Lee JH, Lee J, Choi KY, et al. (2008) Calmodulin dynamically regulates the trafficking of the metabotropic glutamate receptor mGluR5. Proc Natl Acad Sci U S A 105(34): 12575–12580.

Malenka RC and Bear MF (2004) LTP and LTD: an embarrassment of riches. Neuron 44(1): 5–21.

Mander BA, Santhanam S, Saletin JM, et al. (2011) Wake deterioration and sleep restoration of human learning. Curr Biol 21(5): R183–184.

Maris E and Oostenveld R (2007) Nonparametric statistical testing of EEG- and MEG-data. J Neurosci Methods 164(1): 177–190.

McCormick DA and von Krosigk M (1992) Corticothalamic activation modulates thalamic firing through glutamate “metabotropic” receptors. Proc Natl Acad Sci U S A 89(7): 2774–2778.

Mitra A, Snyder AZ, Hacker CD, et al. (2016) Human cortical-hippocampal dialogue in wake and slow-wave sleep. Proc Natl Acad Sci U S A. DOI: 10.1073/pnas.1607289113.

Molle M and Born J (2011) Slow oscillations orchestrating fast oscillations and memory consolidation. Prog Brain Res 193: 93–110.

Mölle M, Marshall L, Gais S, et al. (2002) Grouping of spindle activity during slow oscillations in human non-rapid eye movement sleep. J Neurosci 22(24): 10941–10947.

Mormann F, Fell J, Axmacher N, et al. (2005) Phase/amplitude reset and theta-gamma interaction in the human medial temporal lobe during a continuous word recognition memory task. Hippocampus 15(7): 890–900.

Nettersheim A, Hallschmid M, Born J, et al. (2015) The Role of Sleep in Motor Sequence Consolidation: Stabilization Rather Than Enhancement. J Neurosci 35(17): 6696–6702.

Ngo HV, Martinetz T, Born J, et al. (2013) Auditory closed-loop stimulation of the sleep slow oscillation enhances memory. Neuron 78(3): 545–553.

Niethard N, Ngo HV, Ehrlich I, et al. (2018) Cortical circuit activity underlying sleep slow oscillations and spindles. Proc Natl Acad Sci U S A 115(39): E9220–E9229.

Norimoto H, Makino K, Gao M, et al. (2018) Hippocampal ripples down-regulate synapses. Science 359(6383): 1524–1527.

Oostenveld R, Fries P, Maris E, et al. (2011) FieldTrip: Open source software for advanced analysis of MEG, EEG, and invasive electrophysiological data. Comput Intell Neurosci 2011: 156869.

Pecknold JC, McClure DJ, Appeltauer L, et al. (1982) Treatment of anxiety using fenobam (a nonbenzodiazepine) in a double-blind standard (diazepam) placebo-controlled study. J Clin Psychopharmacol 2(2): 129–133.

Pillai RL and Tipre DN (2016) Metabotropic glutamate receptor 5 – a promising target in drug development and neuroimaging. Eur J Nucl Med Mol Imaging 43(6): 1151–1170.

Porter RH, Jaeschke G, Spooren W, et al. (2005) Fenobam: a clinically validated nonbenzodiazepine anxiolytic is a potent, selective, and noncompetitive mGlu5 receptor antagonist with inverse agonist activity. J Pharmacol Exp Ther 315(2): 711–721.

Rasch B and Born J (2013) About sleep’s role in memory. Physiol Rev 93(2): 681–766.

Rasch B, Born J and Gais S (2006) Combined blockade of cholinergic receptors shifts the brain from stimulus encoding to memory consolidation. J Cogn Neurosci 18(5): 793–802.

Rechtschaffen A and Kales A (1968) A manual of standardized terminology, technique and scoring system for sleep stages of human sleep. Los Angeles Brain Information Service. Brain Information Institute, UCLA.

Sadowski JH, Jones MW and Mellor JR (2016) Sharp-Wave Ripples Orchestrate the Induction of Synaptic Plasticity during Reactivation of Place Cell Firing Patterns in the Hippocampus. Cell Rep 14(8): 1916–1929.

Staresina BP, Bergmann TO, Bonnefond M, et al. (2015) Hierarchical nesting of slow oscillations, spindles and ripples in the human hippocampus during sleep. Nat Neurosci 18(11): 1679–1686.

Steriade M (2006) Grouping of brain rhythms in corticothalamic systems. Neuroscience 137(4): 1087–1106.

Tadavarty R, Rajput PS, Wong JM, et al. (2011) Sleep-deprivation induces changes in GABA(B) and mGlu receptor expression and has consequences for synaptic long-term depression. PLoS One 6(9): e24933.

Tononi G and Cirelli C (2014) Sleep and the price of plasticity: from synaptic and cellular homeostasis to memory consolidation and integration. Neuron 81(1): 12–34.

Walker MP, Brakefield T, Seidman J, et al. (2003) Sleep and the time course of motor skill learning. Learn Mem 10(4): 275–284.

Wen H and Liu Z (2016) Separating Fractal and Oscillatory Components in the Power Spectrum of Neurophysiological Signal. Brain Topogr 29(1): 13–26.

Xu J, Antion MD, Nomura T, et al. (2014) Hippocampal metaplasticity is required for the formation of temporal associative memories. J Neurosci 34(50): 16762–16773.

