## Supplementary Figures for "Specific changes in sleep oscillations after blocking human metabotropic glutamate receptor 5 in the absence of altered memory function"

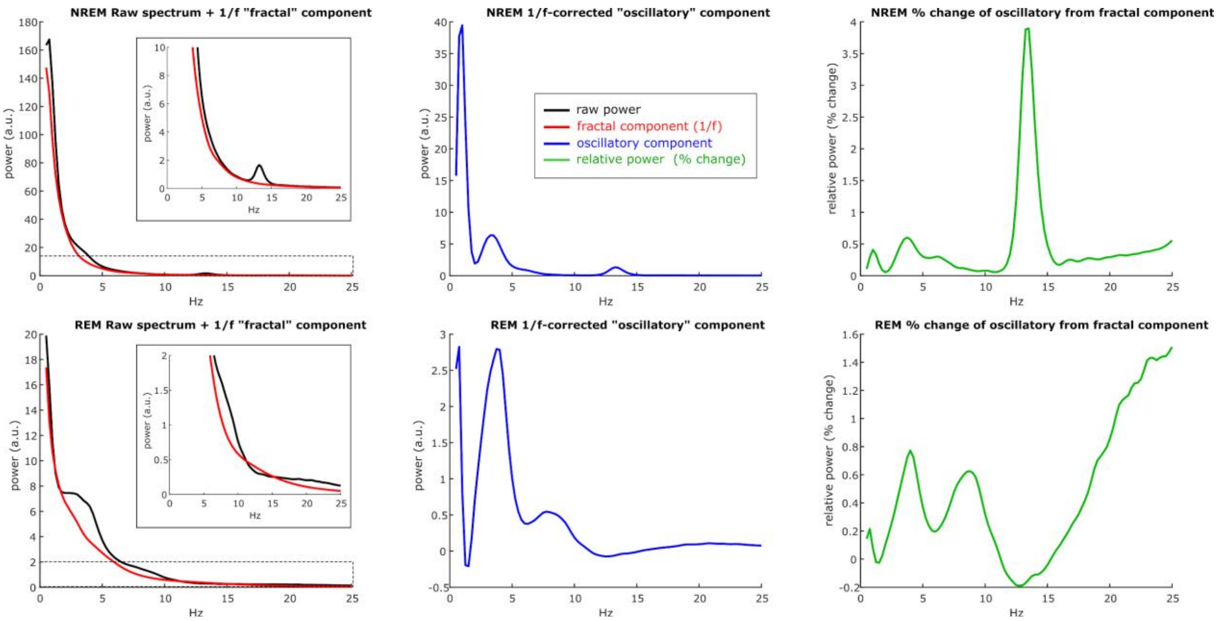


**Supplementary Figure S1**: Example power spectra for Channel Cz of a single subject for NREM (top row) and REM sleep (bottom row), illustrating the processing steps for 1/f-correction and normalization. Subtraction of the “fractal” 1/f-component calculated using IRASA (red) from the original raw power spectrum (black), results in the 1/f-corrected “oscillatory” component (blue). The relative deviation of the “oscillatory” (blue) component from the “fractal” 1/f-component provides the normalized percent change component (green). This procedure removes the common sensitivity bias towards high amplitude low frequency oscillations, allowing all relevant frequency bands to be observed as clear spectral peaks.

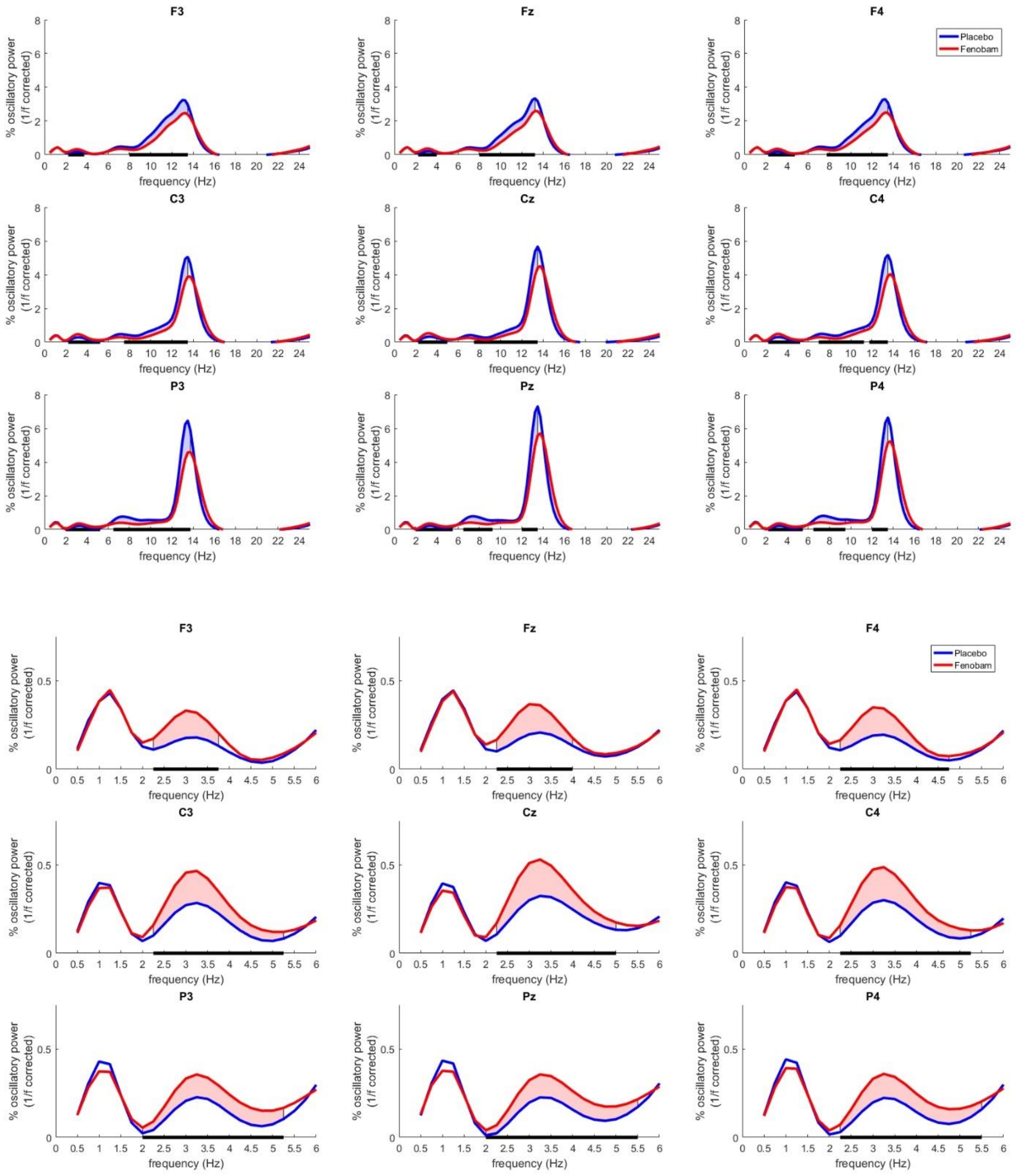


**Supplementary Figure S2:** Frequency domain NonREM sleep.

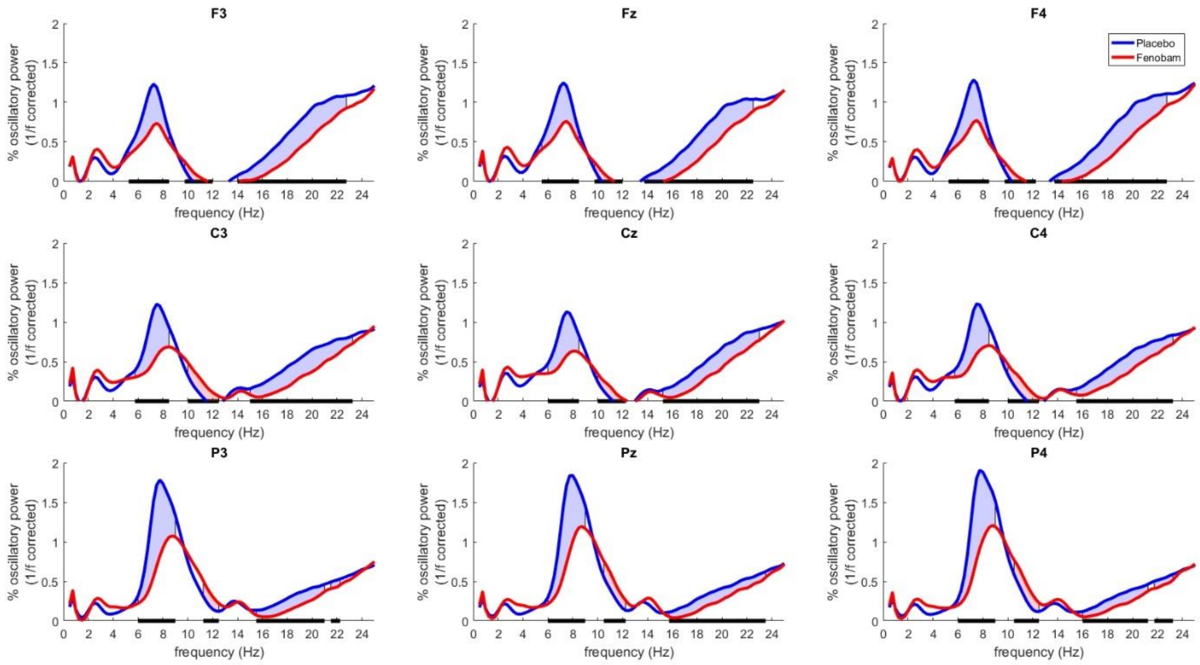


**Supplementary Figure S3:** Frequency domain REM sleep.

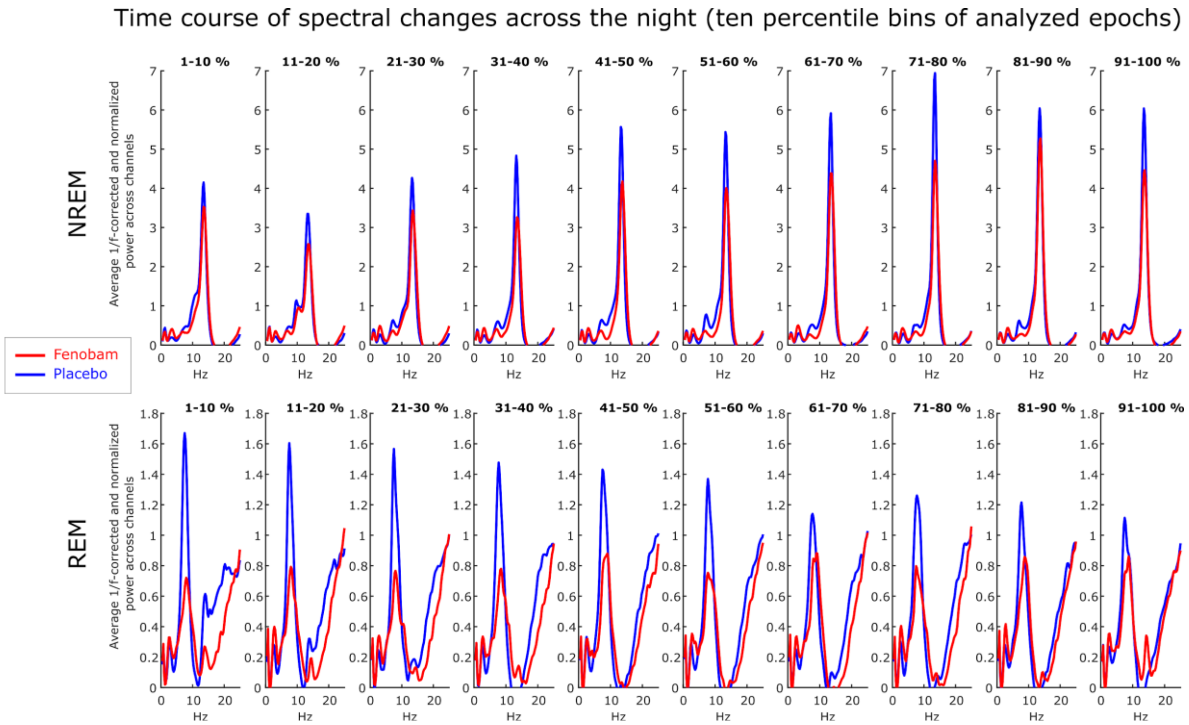


**Supplementary Figure S4**: Power spectra (1/f-corrected and normalized) separately for NREM (upper row) and REM sleep (lower row) and ten consecutive percentile bins of analyzed epochs (columns), with e.g., “1-10%” referring to the first 10% of all artifact-free NREM epochs of each subject, and so forth. Red and blue lines represent Fenobam and Placebo conditions, respectively.

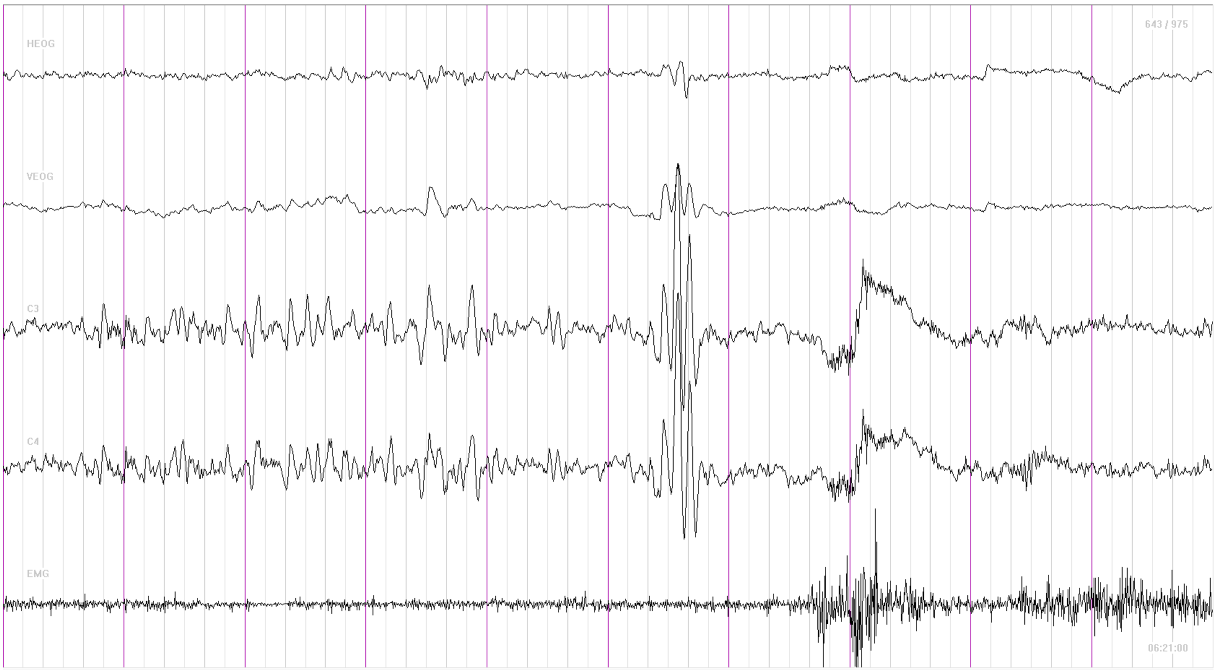


Participant 1 Night 2

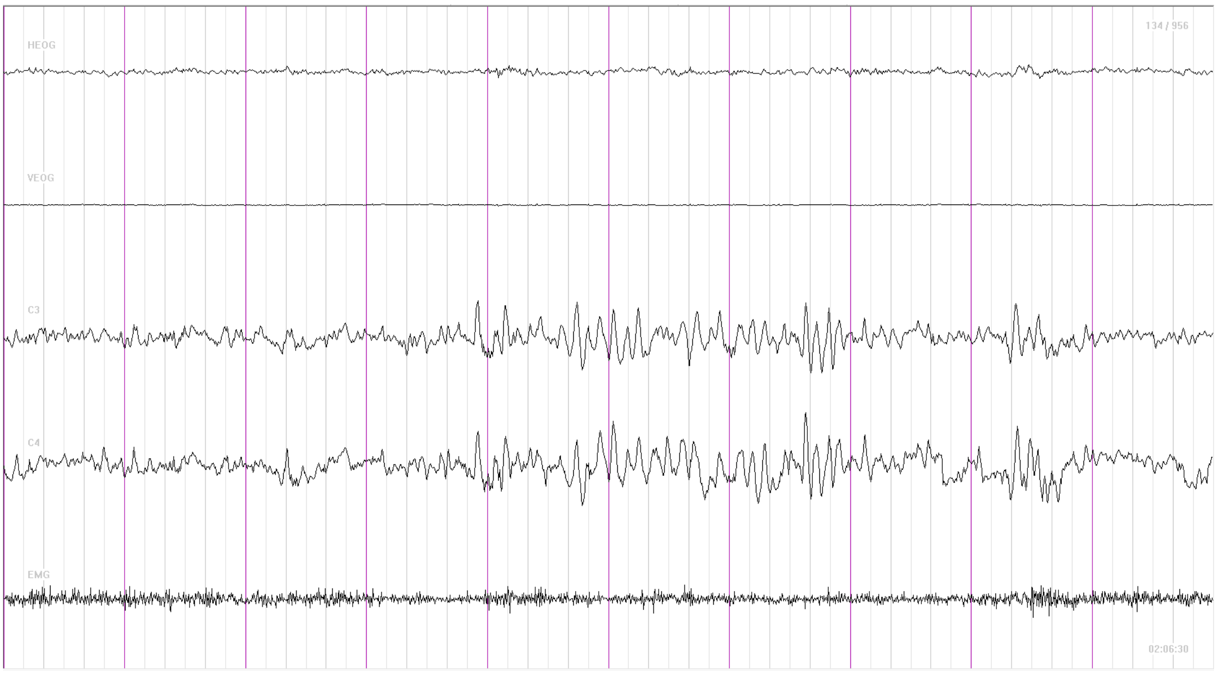


Participant 3 Night 1

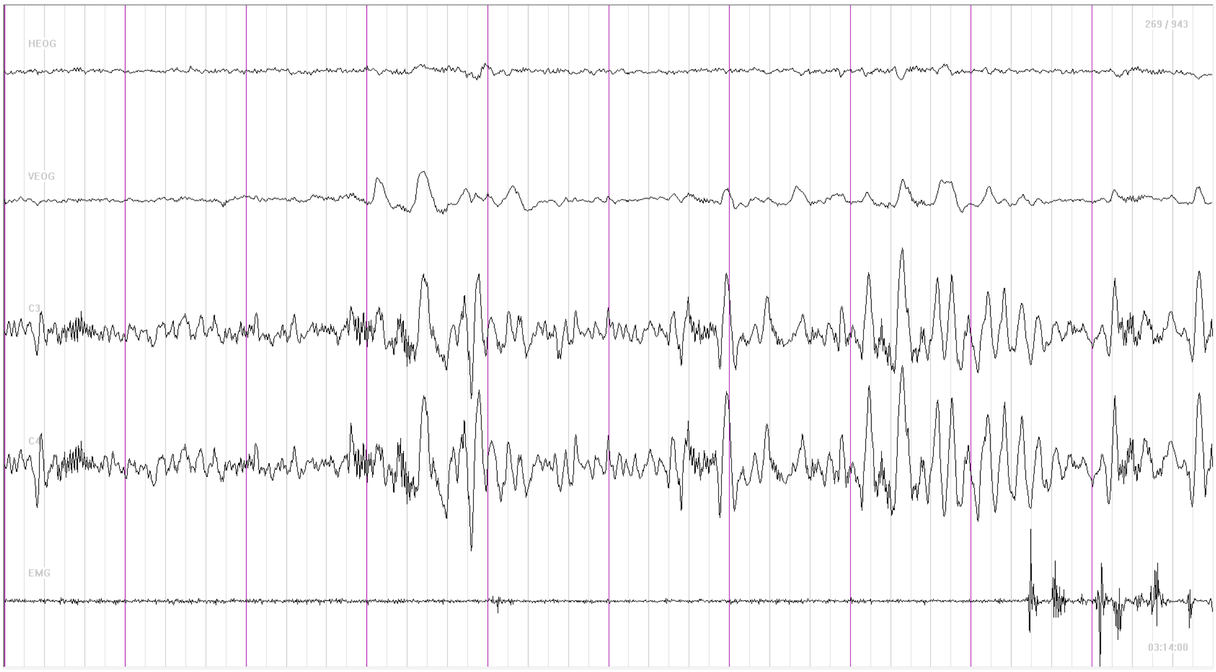


Participant 6 Night 2

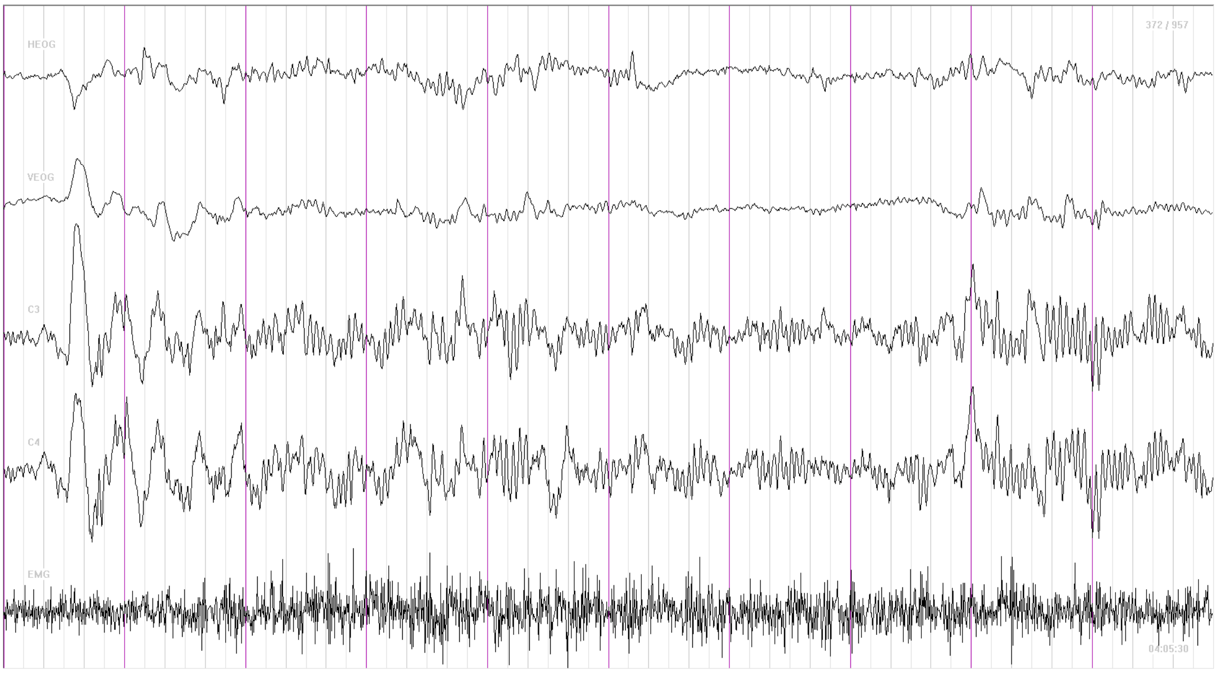


Participant 12 Night 1
